# A network perspective on the role of c-di-GMP-associated protein complexes in biofilm formation

**DOI:** 10.64898/2026.05.12.724550

**Authors:** M-F Noirot-Gros, P Larsen, S Forrester, R Wilton, K. M. Kemner, R. Briandet, G. Babnigg, P. Noirot

## Abstract

The secondary messenger cyclic di-GMP is a ubiquitous bacterial signal that regulates the switch from a free-swimming to a sessile biofilm-forming lifestyle. Many biofilm-forming *Pseudomonas* species possess numerous c-di-GMP-binding proteins (CDGs) which regulate gene expression, protein activity, and protein complexes. However, the mechanisms by which numerous CDG effectors form a coherent signaling network to coordinate lifestyle changes remain poorly understood. We addressed this knowledge gap by focusing on ten CDG proteins involved in biofilm development in *P. fluorescens* SBW25. We used an integrated approach combining a protein interaction network from genome-wide yeast two-hybrid (Y2H) screens with large-scale biofilm and motility phenotype analyses via CRISPR interference (CRISPRi). Our network associated c-di-GMP signaling with processes such as signal transduction, solute transport, secretion, virulence, transcriptional regulation, DNA repair, and cell division. We discovered unknown functions of two CDG proteins in DNA repair and cell division, supporting the significance of our network. Notably, the phosphodiesterase DipA, interacts with numerous CDG proteins through GGDEF domains. Phenotypic analyses revealed that CDG partners were highly correlated or strongly anticorrelated with DipA. These findings suggest that DipA is a central hub for CDG interactions that integrates opposing modules. These findings support the hub-based model of c-di-GMP signaling, which is crucial for localized control and rapid adaptation to environmental changes.

## 2. Introduction

Bacterial biofilms are ubiquitous and represent the dominant microbial lifestyle on Earth^1,2^. Biofilms are structured aggregates of microorganisms encased in a self-produced matrix of exopolysaccharides, proteins, and DNA^1,3,4^. Biofilms exhibit a high degree of resistance to physical and chemical stresses, partly due to the physical barrier provided by the matrix, which enables the bacteria to better withstand nutrient deprivation and external threats^5,6^. They are present in diverse natural environments such as soils, plants, and most terrestrial and aquatic habitats^7^. They are also developing in artificial settings, including industrial water systems, biomedical devices, and food-processing facilities^5,6^. While biofilms are generally regarded as detrimental to human health, they can also confer beneficial effects. Within the rhizosphere, biofilms establish a protective microsystem for plant growth-promoting (PGP) microorganisms by facilitating interactions with plant roots, enhancing nutrient availability, protecting against pathogens, and improving plant resilience to environmental stressors^8,9^.

Biofilm formation is a complex developmental process driven by bacterial sensing and response to environmental cues^10^. This process is largely regulated by the intracellular second messenger cyclic diguanosine monophosphate (c-di-GMP), which is widely found in prokaryotes ^11–13^. C-di-GMP governs physiological and behavioral changes, such as motility, biofilm formation and dispersion, cell differentiation, cell cycle progression, virulence, and quorum sensing^14,15^. In *Pseudomonas* and other bacteria, c-di-GMP plays a central role in the transition from a motile to a sessile biofilm-associated lifestyle by regulating metabolic activity and the production of matrix components such as cellulose and alginate^3,16,17^. Multiple studies have highlighted the role of c-di-GMP in regulating additional metabolic and cellular processes in biofilms^14,15,18–21^. However, the specific metabolic pathways underlying biofilm development and their contributions to this intricate process remain only partially understood.

In bacterial cells, the intracellular level of c-di-GMP is modulated by the balanced activities of biosynthesis via diguanylate cyclases (DGCs) and degradation via phosphodiesterases (PDEs)^13^. DGCs contain a GGDEF signature domain and catalyze the synthesis of c-di-GMP from two GTP molecules. PDEs are characterized by their EAL or HD-GYP domains, and catalyze the degradation of c-di-GMP to the linear dinucleotide pGpG or two GMP molecules ^11^. In addition to their c-di-GMP regulatory domains, PDEs and DGCs often contain additional signaling domains that are generally located at the protein N-terminus. These include HAMP, PAS/PAC, GAF, CACHE, and CHASE, which are involved in sensing extracellular signals^22^. The binding of c-di-GMP to effectors and regulatory proteins consequently impacts cellular functions through various mechanisms including transcriptional, post-transcriptional, post-translational, and protein-protein interactions^23^. The importance of c-di-GMP signaling in promoting biofilm formation is now widely recognized in numerous *Pseudomonas* species, including *P. fluorescens*, which colonizes plant roots, and *P. aeruginosa*, which is responsible for cystic fibrosis and burn wound infections^24–26^. C-di-GMP is suggested to promote the adaptation of pathogenic, but also of beneficial, pseudomonad species to their hosts by shifting their lifestyle from a motile to host-associated biofilm^27,28^. Given the central role of c-di-GMP in bacterial lifetime transition, understanding the regulatory and metabolic pathways controlled by DGCs and PDEs during biofilm formation is important and requires further elucidation.

The c-di-GMP signaling network functions as a regulatory circuit that processes information to facilitate decision-making, such as lifestyle alterations in bacteria. This network is hypothesized to adopt a “bow-tie” architecture integrating many environmental stimuli into intracellular c-di-GMP levels, which subsequently regulate genes associated with biofilm formation or swarming motility^29^. This model explains how signals can be processed through the biochemical computation of many proteins that synthesize, degrade, or bind c-di-GMP to modulate its intracellular concentration on a global scale. However, within bacterial cells, it is postulated that c-di-GMP signaling is facilitated by the integration of both global and localized signaling processes^12,30–32^. These processes coalesce into a complex signaling network, enabling bacteria to convert a wide array of input signals into specific cellular outputs.

Protein-protein interactions involving DGCs, PDEs, and target effectors are essential for establishing localized signaling through the generation of local pools of c-di-GMP^31,32^. Direct interactions between proteins facilitate the assembly of PDEs and DGCs into a signaling complex, serving as a local regulatory node that direct the synthesis or degradation of c-di-GMP, which in turn affects nearby effectors or targets^32^. In *E. coli*, the PDE enzyme YciR, the DGC enzyme YdaM, and the transcriptional regulator MlrA form a c-di-GMP signaling module that regulates the expression of the central biofilm regulator CsgD^33^. Protein-protein interaction studies have demonstrated direct interactions between YdaM, YciR, and MlrA through various domain contacts. Within this signaling complex, YciR acts both as a c-di-GMP effector and a PDE trigger enzyme, degrading locally the c-di-GMP synthesized by YdaM and inhibiting the interaction between YdaM and MlrA by forming a heterodimer with the GGDEF domain of YdaM^33,34^. Another of the best studied decision-making mechanism for biofilm formation controlled by c-di-GMP is the Lap system of *P. fluorescens* Pf01. In this system, the DGC GcbC physically interacts with LapD, a c-di-GMP effector protein, modulating the activity of the protease LapG, which cleaves adhesins, thereby regulating bacterial surface attachment^35^. Further studies on various bacteria have highlighted the role of protein-protein interactions in c-di-GMP-controlled modules^36,37^.

The complexity of c-di-GMP signaling arises from the multitude of DGCs and PDEs encoded in many bacterial genomes. For instance, *P. aeruginosa* and *P. fluorescens* possess about 40 and over 50 c-di-GMP binding proteins (CDGs), respectively^24,30,38^. This complexity is also attributed to the diverse and sophisticated effector systems that translate intracellular c-di-GMP concentrations into specific physiological responses in the host. Key challenges include understanding how cells generate such a wide variety of outputs through a single diffusible second messenger and how bacteria achieve signaling specificity. An increasing body of evidence suggests that protein-protein interactions are central to creating localized signaling events and controlling the complex process of biofilm formation^12,30^. Therefore, the physical interactions between DGCs, PDEs, and their effectors are expected to provide a foundation for deciphering their specific functions, highlighting how bacteria use the c-di-GMP signaling network to make decisions regarding lifestyle changes.

Several cyclic di-GMP (c-di-GMP) signaling proteins (CDGs) are differentially involved in various stages of the *Pseudomonas* biofilm cycle, including surface attachment, maturation, maintenance, and dispersion ^39^. During the initial stage, the transition from swimming to surface attachment is regulated by DGCs such as GcbA, GcbC, WspR, and SadC, along with the PDE enzyme BifA ^40,41^. Similarly, the transition from a sessile to a planktonic state during the dispersion stage involves the DGC GcbA and the PDE DipA ^39,42^. BifA and DipA are dual-domain enzymes containing a GGDEF domain linked to an EAL domain, both functioning strictly as PDEs to degrade c-di-GMP ^42,43^. Despite their common enzymatic activity, BifA and DipA regulate distinct stages of biofilm development and exhibit unique phenotypic characteristics. BifA is involved in the early stage of biofilm formation while DipA is involved during the dispersion stage ^42–45^. In both *P. aeruginosa* and *P. fluorescens* species, the knockout of one gene or the other results in defects in swarming and swimming motility^42,46^. However, a Δ*bifA* mutant exhibits a hyperbiofilm phenotype characterized by a thick and highly structured biofilm, whereas a Δ*dipA* mutant results in a dense but poorly structured biofilm^42,43^.

*P. fluorescens* SBW25 initially isolated from sugar beet, colonizes the roots and enhances plant growth ^47–50^. SBW25 serves as a valuable model organism for investigating biofilm formation dynamics^51–53^. Similar to many *Pseudomonas* species, SWB25 encodes numerous proteins that bind c-di-GMP, which can be categorized based on their protein domains, such as GGDEF, EAL, or other c-di-GMP-binding domains, like PilZ. The biological functions of most of these CDGs remain largely uncharacterized, and their effectors have yet to be identified. Their role in regulating c-di-GMP homeostasis and biofilm formation at the air-liquid interface has been established ^54–56^. The downregulation of *dipA* and *bifA* using CRISPRi-based approaches produced similar biofilm and motility phenotypes as those exhibited by other pseudomonas species ^57^. However, the involvement of c-di-GMP in regulating biofilm and motility phenotypes, particularly in the context of beneficial colonization of plants, requires further investigation.

The yeast two-hybrid approach is a robust technique for identifying protein-protein interactions, offering valuable insights into the biological roles of proteins, including signaling proteins and protein complexes^58,59^. To further elucidate the role of c-di-GMP in regulating bacterial pathways during biofilm formation in *P. fluorescens SBW25*, we conducted a genome-wide yeast two-hybrid screen to identify proteins that physically associate with ten c-di-GMP proteins, which were selected based on their functional roles at various stages of biofilm formation in root-associated *Pseudomonas* species (Table S1). The roles of CDG proteins and their interacting partners in biofilm formation and structure, exopolymeric matrix production, and swarming motility were further investigated using a CRISPR interference (CRISPRi)-mediated phenotyping approach for systematic gene knockdown. Our results revealed a hub-based organization of CDG proteins centered on the phosphodiesterase DipA, as well as numerous connections between CDG proteins with other cellular pathways, including two-component systems, transporters, transcriptional regulators, cell-cell communication, cell division, and DNA repair. We provided experimental evidence supporting the biological relevance of these interactions in DNA repair and cell division, thereby also supporting the significance of this network. Our findings, which include the identification of a CDG hub and its interaction with localized cell processes, are in support of the recently proposed hub-based model for local control of c-di-GMP signaling in the cells^60^.

## 3. Results

### 3.1. A Y2H protein-protein interaction network centered on 10 c-di-GMP binding proteins involved in the formation of biofilm in the rhizosphere

We used the PDEs RimA (PFLU0263), BifA (PFLU4858), DipA (PFLU0458), RbdA (PFLU4308), RapA-like (PFLU2031), and putative PDE PFLU1114, the DGCs PFLU5127, AdrA (PFLU3650), and AswR (PFLU5210), and the PilZ motif containing protein Alg44 (PFLU0988) as baits in genome-wide yeast two-hybrid screens. These c-di-GMP binding baits were selected based on i) their involvement in different steps of biofilm formation and dispersal as reported in the literature, and ii) their involvement in adapting to environmental changes such as during colonization of plant roots, as explained in greater detail in Supplementary information SI-1 and in Table S1. Notably, a specific design was made for AdrA to exclude the N-terminal multipass membrane domain. Our analysis identified 99 independent high-confidence interactions and 127 moderate-to-low confidence interactions (Table S2). To assess the reliability of each interaction, a Predicted Biological Score (PBS) was computed as previously described^61,62^. This score provides confidence levels ranging from high biological relevance (category A) to low probability (category D). The PBS serves as a statistical confidence measure, indicating the likelihood that a detected protein–protein interaction is biologically authentic rather than an experimental artifact. It achieves this by integrating a local score, which considers the number and independence of the prey fragments recovered and their reading frames, with a global score that estimates the probability of a nonspecific interaction based on the frequency with which each prey is identified across all other screens conducted with the same library^61,62^. Interaction partners were classified as ‘high-confidence’ if they were repeatedly and independently identified in the same screening. Consequently, a high-confidence network map was constructed, linking 94 distinct proteins through 110 highly significant interactions (Fig. 1A). Approximately 78% of the identified protein partners were detected multiple times as independent and overlapping fragments, outlining the distinct minimal domain involved in the interaction (Table S2). This protein-protein interaction (PPI) network, centered on the c-di-GMP signaling pathway, unveiled protein complexes likely to have biological functions within the cell. Within the network, the partner proteins were categorized into various functional groups, linking the c-di-GMP pathway to many cellular processes, including nutrient transport, secretion, virulence, phosphorylation, transcriptional regulation, and DNA repair (Table S3). The distribution of functional annotations among the interacting partners of each c-di-GMP bait protein was analyzed to determine the statistically significant enrichment of annotations (Fig. 1B, Table S3). Among the most significantly enriched functional categories, the signal transduction pathway was predominantly associated with AwsR (PFLU5210) and Alg44 (PFLU0988), whereas the c-di-GMP signaling pathway was the most highly represented for DipA (PFLU0458) and PFLU1114. The DNA transactions annotation was strongly linked to AdrA (PFLU3650) and RimA (PFLU0263) while transcription was significantly connected to BifA (PFLU4858) and PFLU2031. RbdA (PFLU4308) and PFLU5127 were significantly enriched in protein partners in the adhesion and toxicity categories, and AdrA was also associated with redox activity (Fig. S1A, Table S3). These findings align with research conducted on various bacterial species, which has demonstrated that c-di-GMP is intricately linked to multiple cellular pathways and regulatory mechanisms ^63^.

**Figure 1:**
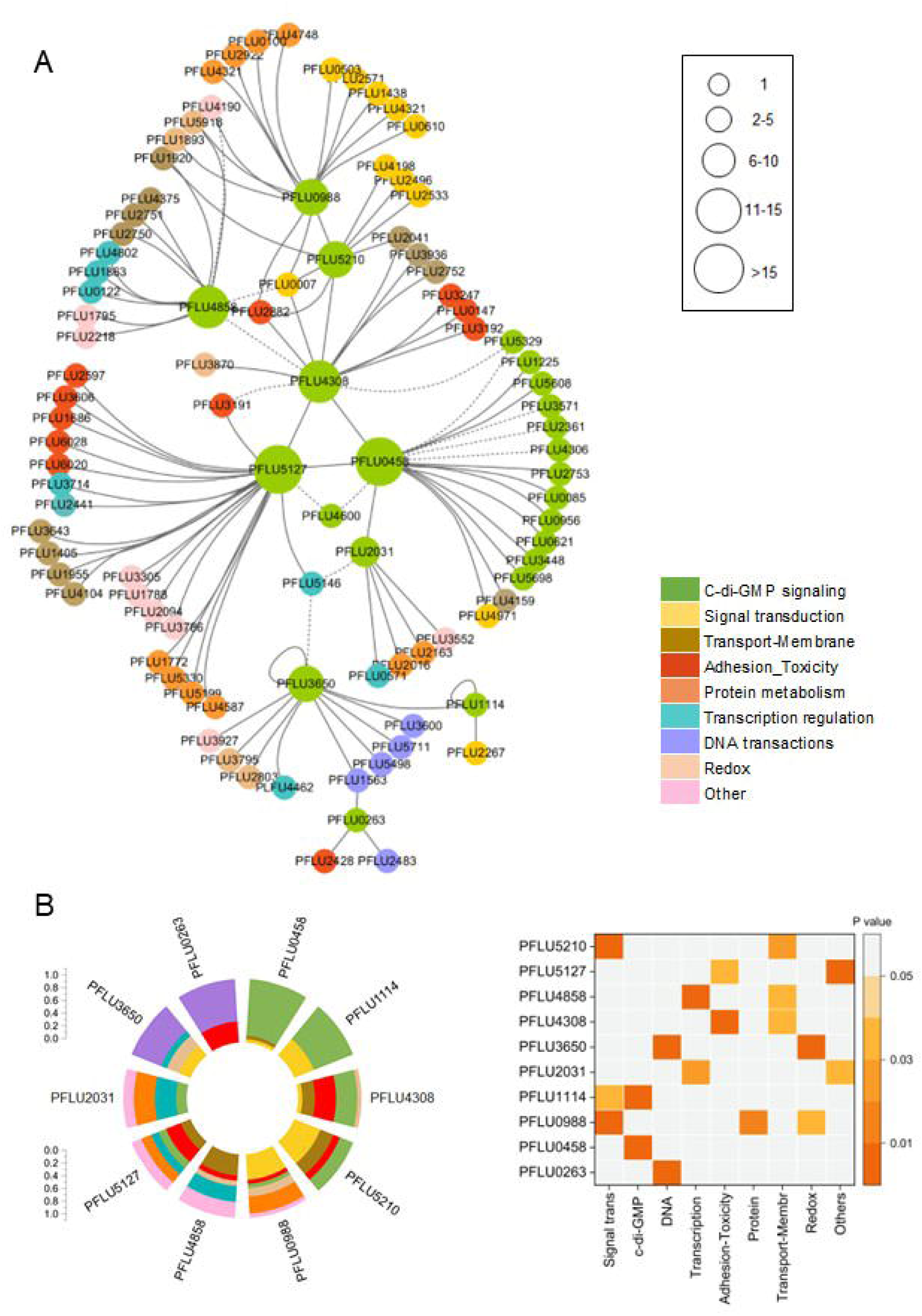
Y2H protein-protein interaction network centered on c-di-GMP signaling. A. Proteins are represented as nodes colored according to their functional annotations, and edges indicate interactions between nodes with good-to-high (solid line) and moderate (dotted line) confidence, as described in Table S2. B. Left: Distribution of the functional categories of the interacting partners for the ten c-di-GMP bait proteins used in this study (extracted from Table S3). Right: Statistical significance of functional category enrichment for each c-di-GMP bait protein, calculated as cumulative hypergeometric distribution (from data in Table S3). P-values less than 0.05 are highlighted in color.

#### 3.1.1. A DipA-centered c-di-GMP signaling network

Within the PPI network, the initial set of 10 c-di-GMP bait proteins was augmented by 23 c-di-GMP binding proteins, constituting approximately half of all CDG proteins documented in SBW25. Notably, 16 c-di-GMP-binding proteins interacted with DipA (PFLU0458). Among these DipA-associated proteins are DgcP (PFLU0085), WspR (PFLU1225), GcbA (PFLU0621), GcbB (PFLU4600), GcbC (PFLU0956), MorA (PFLU5329), RapA (PFLU2031) and MucR (PFLU2753) (Fig. 1A, Fig. S1A, Tables S2 and S3). Furthermore, DipA was identified as a binding partner of RbdA (PFLU4308), RapA (PFLU2031), and PFLU5127 when employed as baits in Y2HB genomic screens. Consequently, the prevalence of CDG proteins associated with DipA is very high, representing 25% of all proteins in the network. This suggests that DipA may function as a CDG signaling hub protein in the cell. Supporting this notion, a detailed examination of the interacting fragments of the DipA CDG partners revealed the presence of the GGDEF regulatory domain within the minimal domains necessary for interaction with DipA (Fig. 2A). This finding suggests a singular and common mode of interaction between DipA and all its CDG partners, potentially through heterodimerization of their respective GGDEF domains. In addition to DipA, the PDE RbdA (PFLU4308) was identified as a prey protein for BifA (PFLU4858), AswR (PFLU5210), and PFLU5127 within the network (Fig.1A, Fig.S1-A, B). The RbdA-interacting fragments included the PAS/PAC signaling domain, either exclusively (for BifA and AwsR) or in addition to the GGDEF domain (for PFLU5127 and DipA) (Fig. S1-B). This observation suggests that CDG proteins may also engage with other CDG partners using different functional domains.

**Figure 2:**
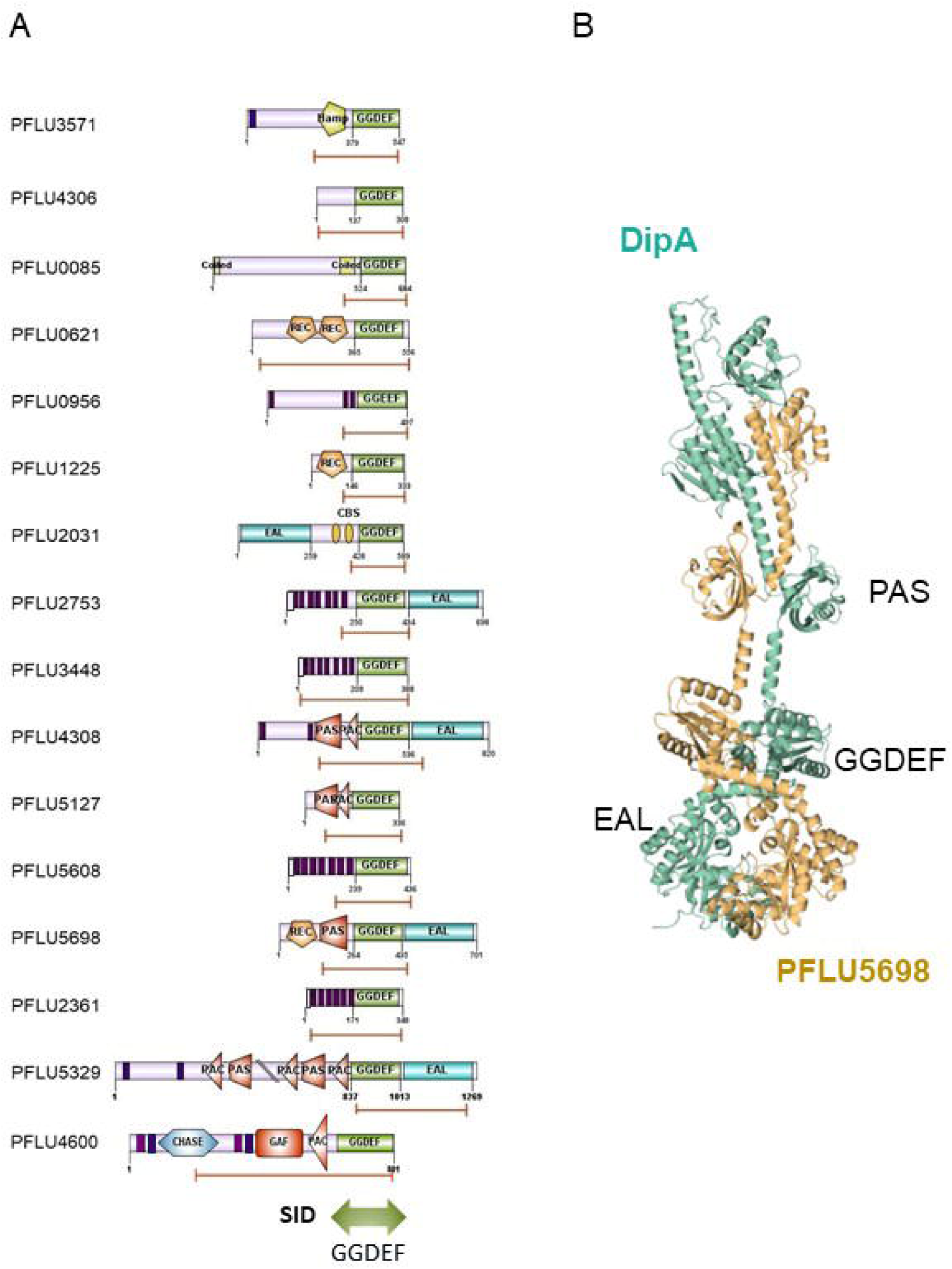
Delineation of the minimal interacting domain. A- The functional domains of all DipA (PFLU0458) partners are depicted as predicted by SMART (https://smart.embl.de/). GGDEF and EAL domains are shown in green and light blue rectangles, respectively. Accessory domains are indicated in orange pentagon (REC), green pentagon (HAMP), blue hexagon (CHASE), red rectangle (GAF), red triangle (PAC), red pentagon (PAS), and dark violet segments (transmembrane). For each DipA interacting partner, the minimal interacting domain was indicating by capped brown lines. The minimal interacting domain with DipA spans the GGDEF domains of all partners. B-Structural model of the best ranking DipA-PFLU5698 complex as predicted by Boltz-2 (Supplementary information SI-2). DipA (light green); PFL5698 (light orange) with EAL, GGDEF and PAS domains as indicated.

#### 3.1.2. Crosstalk between c-di-GMP and phosphorylation signaling pathways

The second largest functional category identified in the network comprised two-component signal transduction systems (TCS), including methyl chemotaxis systems (MCS). These were notably enriched among the partners of Alg44 (PFLU0988) and AswR (PFLU5210), underscoring the propensity of these two CDG proteins to associate with components of phosphorylation-based signaling pathways (Fig. 1B, Fig. S1A, Tables S2 and S3). A compelling example is the transmembrane sensor histidine kinase PFLU0007, which interacts with Alg44, AswR, RbdA, and BifA (Fig. S1-C). PFLU0007 consists of an N-terminal extracellular sensory domain (CHASE) linked to a cytoplasmic C-terminal signaling domain. The C-terminal signaling domain contains a PAS module and a HisKA/HATPase module, which is responsible for histidine phosphotransfer and homodimerisation^64,65^. The fragments of the PFLU0007 protein identified as interacting with the four CDG proteins were mapped to the cytoplasmic C-terminal half, encompassing the PAS and HisKA/HATPase domains (Fig. S1C). PFLU0007 and its four partners formed an interconnected cluster, suggesting a complex interplay between c-di-GMP and phosphorylation signaling pathways in SBW25. Collectively, the two functional categories, c-di-GMP signaling and signal transduction, predominated, accounting for approximately half of the interactions within the network (Fig. 1B).

#### 3.1.3. c-di-GMP and transmembrane transport

Our dataset also revealed a high proportion of proteins involved in the uptake and efflux of components across the membrane (Fig. 1A, Tables S2 and S3). This functional category was statistically highly enriched in the BifA (PFLU4858) interactome and was also prevalent among RbdA (PFLU4308) and PFLU5127 partners (Fig S1A, Fig.2C). Notably, BifA interacts with MdtB (PFLU2750) and MdtC (PFLU2751), two subunits of the RND inner membrane multidrug transporter MdtABC, as well as with PFLU2720 and PFLU3215, which are homologous to the other RND transporters AcrD and AcrB, respectively. These RND transporters are responsible for expelling many compounds, including enterobactin siderophore^66^. Additionally, CusA (PFLU3936) and CusC (PFLU2752), which encode components of the RND-type Cu/Ag efflux pump responsible for cell detoxification, interacted with RbdA. Finally, the network revealed that PFLU2041, a metal ABC transporter substrate-binding protein (SBP), interacts with RbdA (PFLU4308) and AswR (PFLU5210) with a very high degree of confidence. Indeed, 14 independent fragments of PFLU2041 covering the 80-amino acid N-terminal domain of PFLU2041 were identified in the screening with RbdA (Tables S2 and S3). The PFLU2041 80 pb N-terminal fragment also bound to AswR (Tables S2 and S3). This observation suggests the potential involvement of two transmembrane CDGs, RbdA and AwsR, which possess phosphodiesterase and diguanylate cyclase activity, respectively, in regulating extracellular metal transport^67^.

#### 3.1.4. Toxin delivery and adhesion

CDG proteins associated with two-partner secretion (TPS) toxin-antitoxin (TA) systems represent another significant functional category within this network. TPS is involved in the secretion of cytotoxic compounds or virulence factors and plays a role in pathogenesis, host interaction, and interbacterial competition^68^. This functional category was significantly enriched in the RbdA (PFLU4308) and PFLU5127 interactomes and was also linked to RimA (PFLU0263) (Fig. 1B, Fig. S1A, Tables S2 and S3). Rearrangement hotspot systems (RHS) are TPS delivery systems involved in the contact-dependent growth inhibition (CDI) of neighboring cells^69,70^. PFLU5127 exhibited high-confidence interactions with all four RHS proteins identified in the network: PFLU1686, PFLU2597, PFLU3606, and PFLU6028. The minimal domains necessary for interaction with PFLU5127 were consistently located in the C-terminal region of the RHS shell, excluding the terminal toxin domain, suggesting a similar binding mode to that of PFLU5127. RbdA interacted with three proteins annotated as adhesin activators of the HlyB/ShlB family: PFLU3192, PFLU0147, and PFLU3247. In addition, PFLU3191, an adhesin part of a putative TPS with PFLU3192, was identified to bind PFLU5127 (Fig. S1A, Tables S2 and S3). These observations imply that TPS may be regulated by the c-di-AGMP signaling pathway. Notably, in the PPI-network, the Phage-tail Tape Measure Protein (TMP) PFLU2882 was directly connected to four CDG proteins: AswR (PFLU5210), RbdA (PFLU4308), BifA (PFLU4858), and Alg44 (PFLU0988). TMP proteins, originally described as phage-tail length control proteins, also belong to a subclass of bacterial type VI secretion systems (T6SS) resembling a phage tail, designed to inject toxins directly into neighboring bacteria^69,71,72^. In *P. aeruginosa*, TPM proteins have been shown to promote bacterial cell lysis, leading to the release of eDNA, which is important for forming the biofilm matrix. These observations suggest that these bacterial defense and killing machineries are regulated by cyclic di-GMP.

#### 3.1.5. Transcriptional regulation and DNA repair

Our network also included several transcriptional regulators of diverse families, as well as the heat shock sigma factor σ24 associated with BifA (PFLU4858), RapA (PFLU2031), AdrA (PFLU3650), and PFLU5327 (Fig.1, Fig. S1A, Tables S2 and S3). This functional category was significantly enriched for the BifA and RapA bait proteins (Fig.1B). Three transcription regulators, PFLU5146, PFLU2441, and PFLU3714 were connected to PFLU5127 with a high confidence, suggesting the potential involvement of this putative diguanylate cyclase in modulating their function. Finally, the network revealed a connection between multiple proteins belonging to the DNA repair pathway with the DGC AdrA (PFLU3650) and PDE RimA (PFLU0263) (Fig. 1, Fig. S1A, Table S2 and S3). Notably, AdrA formed high-Aconfidence interactions with PFLU5498 and PFLU3600, two proteins homologous to subunit A of the UvrABC complex, which is involved in the nucleotide excision repair pathway^73^. In addition, AdrA interacts with DNA ligase LigB (PFLU5711), which is involved in oxidative DNA damage repair ^74^. Furthermore, both AdrA and RimA were associated with PFLU1563, an annotated DNA helicase.

### 3.2. Benchmarking protein complex structure prediction against the yeast two-hybrid interactome of DipA

To assess the ability of AI-based 3D structure model to predict our experimentally determined protein-protein complexes, we conducted an analysis of the DipA (PFLU0458) interaction dataset (see Supplementary Information SI-2), using the Boltz-2 structural biology foundation model ^75^ Predicted interactions were benchmarked against three annotated reference sets, which included Y2H-positive interactors to DipA (n=14), putative negative interactors possessing a GGDEF domain (n=3) and sequence-distant proteins with detectable sequence similarity to PFLU0458 (n = 3). Importantly, proteins from the last two categories were not identified as interactors to DipA in our Y2H screen. Protein complexes were predicted with Boltz-2 via the Neurosnap platform and evaluated using the structure-based binding affinity predictor PRODIGY. Each bait and prey sequence pair were modeled as a 1:1 heterodimer, and five independently seeded models were generated. Confidence metrics and interface score were derived from each Boltz-2 prediction and the best (rank1), maximum, mean, and standard deviation values were calculated across the five models (see Supplementary Information SI-2). We found that Boltz-2 did not predict a confident, contact-rich interface for any of the tested complex and did not distinguish Y2H-positive interactions from the control datasets (Supplementary information SI-2). Among the fourteen Y2H-positive interactions, the PFLU0458–PFLU5698 complex was the best ranked model based on interface score parameters and predicted binding free energy using PRODIGY, achieving a moderate range ipSAE score of 0.41 with approximately 451 intermolecular contacts spanning the GGDEF domain and a mean binding free energy of ΔG = −37.4 ± 3.1 kcal·mol⁻¹ (Table 2 in Supplementary information SI-2). This putative 3D structure model is represented in Fig.2B. However, comparison of predicted binding affinities for the best scoring complexes in each of the 3 groups (Y2H positive, putative negative, sequence distant) revealed that binding affinities correlated with the number of predicted intermolecular contacts rather than with the Y2H positive status (Table 3 in Supplementary information SI-2). Altogether, these findings suggest that current structure-based confidence metrics and subsequent binding affinity estimations are insufficient for identifying transient signaling interactions within our c-di-GMP regulatory networks. This finding is consistent with the reported challenges of predicting transient protein–protein interactions within signaling pathways, which may be influenced by allosteric modulation by intracellular second messengers such as the c-di-GMP or by post-translational modifications ^76^. Generally, the standard confidence thresholds developed for stable protein complexes may not be directly applicable to this class of interactions.

### 3.3. CRISPRi-based silencing of c-di-GMP binding proteins and their protein partners impact biofilm and motility phenotypes

We employed a CRISPRi-based approach to systematically suppress the expression of genes encoding network proteins, aiming to elucidate the role of CDGs and protein partners in the motile and sessile behaviors of bacteria (Fig. 3A, Table S4). Our CRISPRi knockdown system has been previously utilized for the phenotypic analysis of biofilm genes in SBW25, demonstrating its ability to produce phenotypes akin to those of gene-knockout strains^77^. In this study, we focused on the downregulation of 23 genes encoding all the CDG proteins that were present in the network and encompassing all the DipA CDG protein partners. Additionally, we selected a group of 33 genes encoding interacting partners and comprising representatives of each functional category identified for each CDG bait, with the aim to provide a more comprehensive overview of the diverse functions potentially involved in biofilm formation (Fig. 3A, Tables S2 and S3). We included a third out-of-network group composed of 14 genes encoding CDG proteins that were absent from our network, to be able to assess the specificity of in-network CDG toward biofilm formation (Fig. 3A). All strains, including a control strain expressing no guide RNA, were grown under the same conditions in the presence of anhydrotetracycline (aTc) to induce downregulation and were incubated at 25°C with 70% controlled humidity for 48 h prior to phenotypic observation (see Materials and Methods). Swarming motility was assessed by determining the swarm area of the strains on semi-solid surface media (Fig.3B and Fig. S2). Air-liquid biofilm pellicles were observed using spinning disk confocal microscopy (SDCM), and parameters related to biomass (biovolume BV, biomass BM), thickness (maximum height MX, mean thickness MS), and roughness (R) were extracted from image analysis using the biofilm extension package of the IMARIS software (Fig.3C and Fig. S3). To assess the capacity to form a submerged surface-associated biofilm, cells adhering to the surface of the pegs of an MBEC™ biofilm incubator immersed in a liquid medium were enumerated by flow cytometry (submerged biofilm at pegs, SBP, Fig. 3D). Given the viscous nature of the biofilm matrix formed by SBW25, this parameter was deemed suitable for measuring increased biofilm levels. However, it was found to be less reliable when measuring low quantities of biofilm material on the pegs (see Materials and Methods). Finally, we utilized a non-toxic red fluorescent optotracer (EbbaBiolight 630) to monitor the production of curli amyloid fibers (AMF) and cellulose in extracellular matrix polymers in growing cultures (Fig.3E, Fig. S4). For each strain with a silenced gene, the phenotypic parameters were standardized to the control strain in the same experiment, consisting of SWB25 expressing *dcas9* but lacking a targeting guide RNA. All the phenotypic data are presented in Table S5.

**Figure 3.**
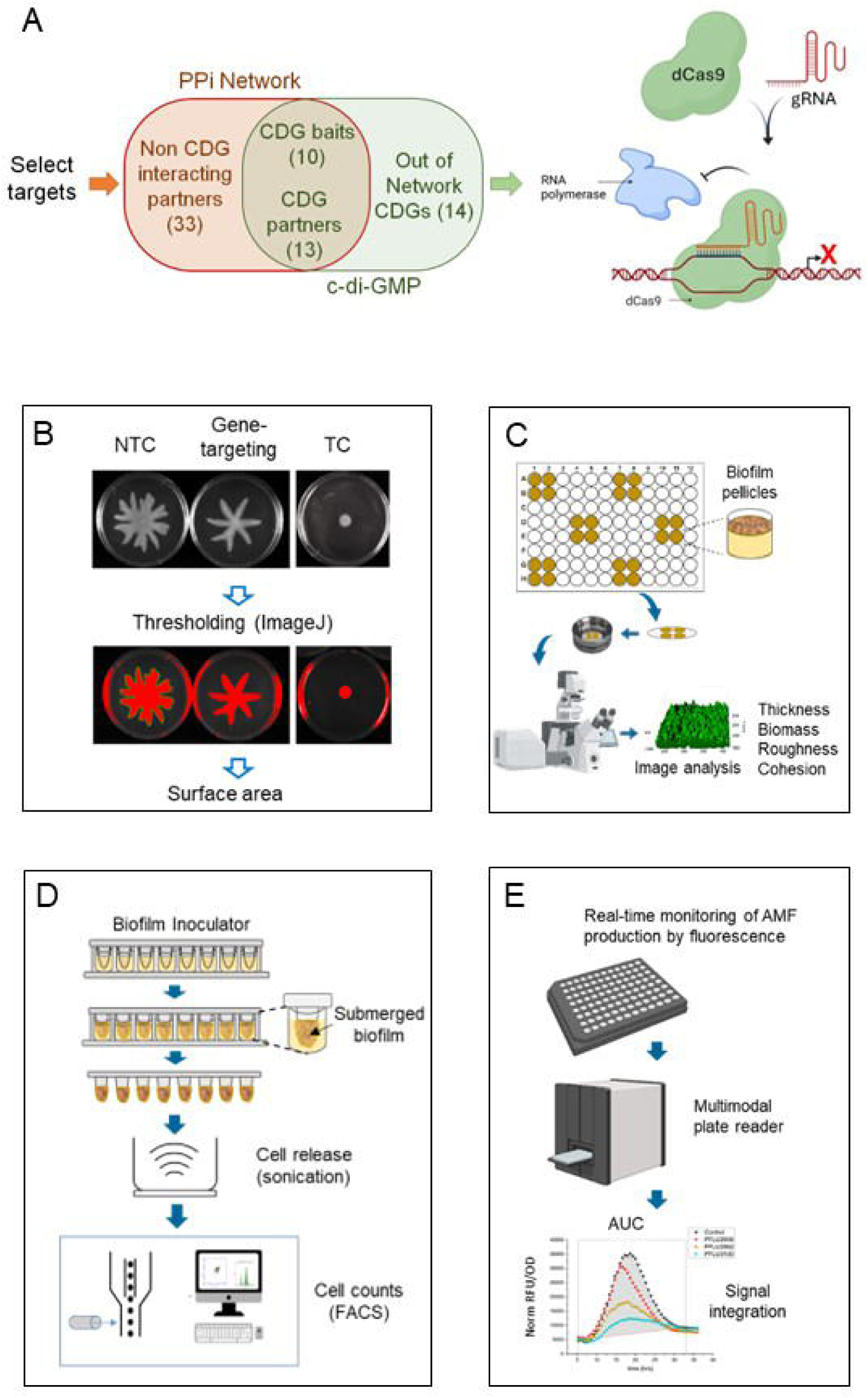
CRISPRi-mediated biofilm and motility phenotyping. A- Specific set of genes encoding in-network CDG proteins, their interacting partners, and out-of-network CDGs were targeted for silencing using a dCas9/gRNA complex through CRISPR interference (CRISPRi), and analyzed for their effects on various biofilm and motility phenotypes. B- Swarming assay, imaging, and calculation of swarm areas. NTC, non-targeting control; TC, targeted positive control. C- Biofilm pellicle assay and imaging by confocal microscopy. Pellicles at the liquid-air interface were peeled out onto circular glass coverslips and mounted on a microscopy chamber containing transparent media. Biofilms were stained with FilmTracer^TM^ green fluorescent dye prior to volume imaging using confocal microscopy. Parameters were extracted by image analysis using Imaris software. D- Formation of submerged biofilm at pegs (MBEC^TM^). Cells were released from the pegs by sonication and counted using flow cytometry. E- Real-time monitoring of amyloid fibers and extracellular matrix production in the presence of optotracer using fluorometry (EbbaBiolight680).

#### 3.3.1. Phenotypic profile of the network

To investigate the organization of phenotypic responses, we conducted hierarchical clustering of standardized Log_2_fold changes across all traits related to biofilm and motility (swarming). This analysis identified distinct clusters of gene knockdowns exhibiting similar phenotypic signatures across different phenotypes (Fig. 4A). Three main phenotypic categories emerged. The first category included genes whose silencing resulted in a reduction in biofilm volume and mass (Fig.4A, red cluster). The second, larger category was characterized by a decrease in swarming ability and an increase in submerged biofilm production following knockdown (Fig. 4A, yellow cluster). The third category comprised genes whose silencing led to a reduction in the production of amyloid matrix components (Fig. 4A, light blue cluster). Overall, these findings underscore that silencing of most genes from the network exhibited at least one biofilm or swarming phenotype, indicating that our network is highly enriched in proteins with biofilm- and motility-related functions.

**Figure 4:**
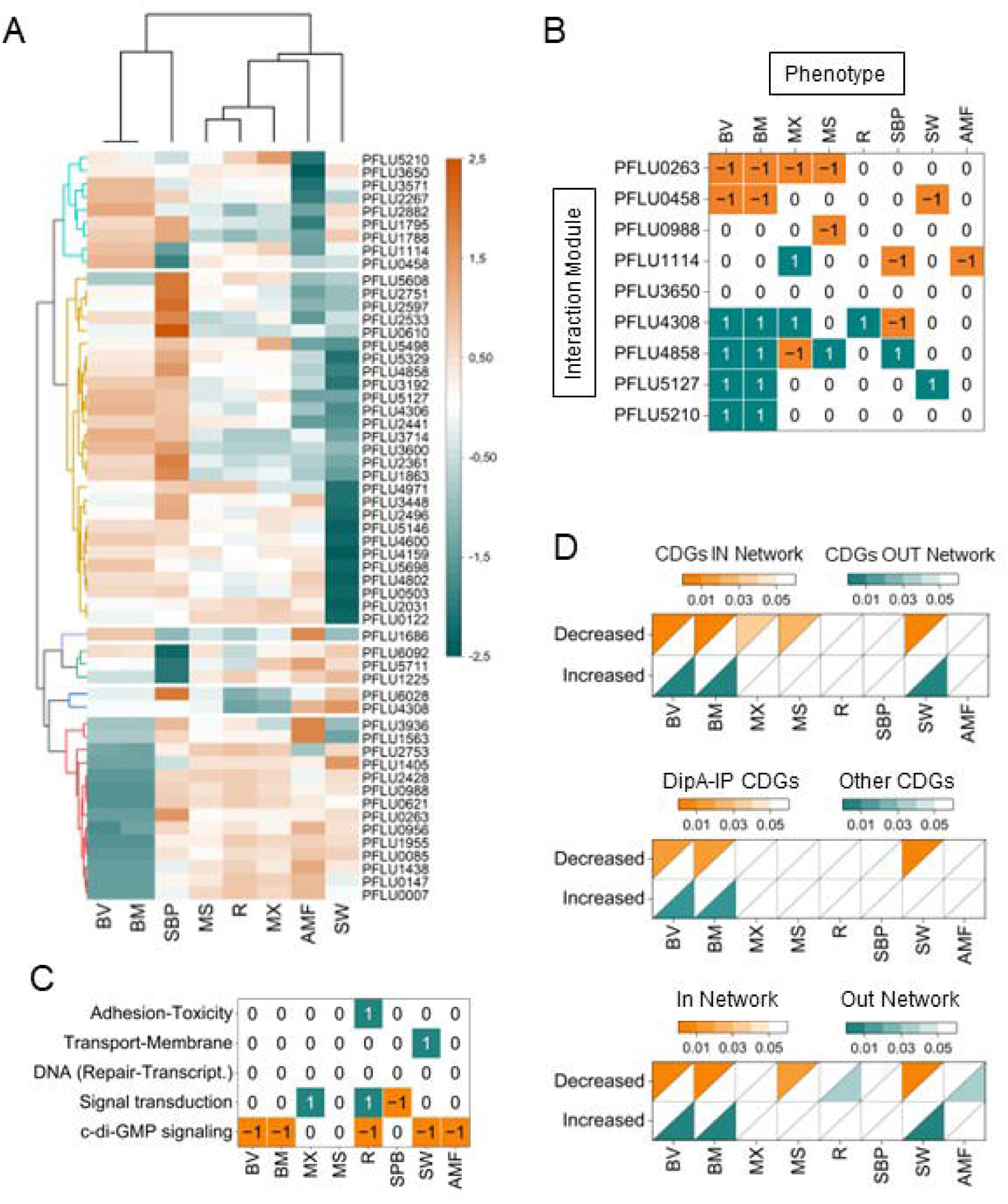
Phenotypic analysis. A- Heatmap with dendrogram of hierarchical clustering of network proteins according to their phenotypic traits. The analyzed phenotypes were biofilm biovolume (BV), biomass (BM) mean thickness (MS), maximum height (MX), roughness (R), submerged biofilm at pegs (SBP), and production of amyloids (AMF), and swarming motility (SW). The targeted proteins (rows on the heatmap) are split into six groups by k-means clustering. Groups are indicated by colors (left side) and distance is Pearson correlation. The extent of phenotype change is indicated by the side bar, green-blue for decrease and orange for increase. (B-D)- *Statistical analysis of phenotypic enrichment.* The analyzed phenotypes were biofilm biovolume (BV), biomass (BM), mean thickness (MS), maximum height (MX), roughness (R), submerged biofilm at pegs (SBP), production of amyloids (AMF), and swarming motility (SW). Statistical significance was calculated using cumulative hypergeometric distributions. All P-values are reported in Table S6. B- Phenotypic enrichment within interaction modules comprising a CDG bait and its direct interaction partners. 1 = significant enrichment for gene partners that increase the phenotype; −1 = significant enrichment for gene partners that decrease the phenotype; and 0 = no enrichment. Note that a value of ‘no enrichment’ does not indicate a lack of change in the phenotype, rather it indicates that the observed frequency of change is equivalent to random chance given the distribution of phenotypes across all knockdowns. C- Phenotypic enrichment by annotation category. D- Phenotypic effects of in-network versus out-of-network knockdowns. Split heatmaps illustrating -Top: phenotypic decrease or increase for knockdown of genes encoding CDG proteins from the PPI network (CDGs IN Network, orange) versus the CDGs not present in the network (CDGs OUT-Network, blue); - Middle: Gene knockdowns of DipA interacting partners (DipA-IP CDGs, orange) versus all other CDGs (blue); - Bottom: Knockdown of all genes present in the network (IN Network, orange) versus those outside of the network (OUT Network, blue).

#### 3.3.2. Phenotype enrichment analysis

We investigated whether the CDG modules, comprising a CDG bait protein and its interaction partners, were enriched in genes displaying one or more specific biofilm and motility phenotypes, using our high-confidence PPI and phenotypic data with statistical significance (Fig. 4B, Table S6). Our findings revealed that the knockdown of genes in the DipA (PFLU0458) interaction module exerted a significant negative impact on biofilm mass and swarm size. Given that all the DipA-binding partners in the network were targeted by CRISPRi, this inhibition emerged as our most significant observation, corroborating the involvement of DipA and its interacting partners in biofilm formation and motility. Similarly, knockdown of the RimA (PFLU0263) interaction subnetwork significantly decreased biofilm mass, volume, and thickness, indicating a positive regulation of biofilm formation. The knockdown of Alg44 (PFLU0988) and its partners is more likely to result in a decrease in biofilm thickness, as already observed^77^. Although AdrA (PFLU3650) subnetworks did not exhibit significant enrichment of phenotypes, the knockdown of genes within the RbdA (PFLU4308), BifA (PFLU4858), PFLU5127, and AwsR (PFLU5210) interaction modules revealed an enrichment of enhanced biofilm phenotypes, suggesting an inhibitory role in biofilm formation. Finally, the knockdown of PFLU1114 and its partners had a positive effect on surface thickness and a negative effect on submerged biofilm and amyloid fiber formation, suggesting a role in biofilm structure.

Next, we examined whether phenotypic changes correlated with the functional categories of the 56 network genes subjected to knockdown (Fig. 4C). Using categorical functional annotations (Table S3) and biofilm and motility phenotypes upon knockdown (Table S5), we found that c-di-GMP signaling genes were significantly enriched in decreased biofilm phenotypes, including biovolume (BV) and biomass (BM), roughness (R), and secretion of amyloid components of the matrix (AMF), as well as decreased swarming ability (Fig. 4C, Table S6). These findings were anticipated and are aligned with the established role of the c-di-GMP pathway in biofilm formation, structure, and motility. The downregulation of genes within the signal transduction category was significantly enriched for enhanced roughness and increased maximum pellicle height, both phenotypes being associated with biofilm structure. These results suggest that protein phosphorylation may play a role in regulating biofilm structural properties. Silencing genes from the transport and membrane classes was significantly associated with an increased swarming phenotype, suggesting that these genes may negatively modulate the swarming behavior. Additionally, silencing genes from the “adhesion-toxicity” category significantly decreased biofilm roughness. Finally, silencing genes involved in transcription regulation and DNA repair did not enrich any of the tested biofilm and motility phenotypes, indicating that the frequency of change was not statistically significant. Our results confirm the established functional relationship between the c-di-GMP signaling pathway and biofilm and motility phenotypes, while also revealing potential functional links with other cell processes, including adhesion and toxicity, transcription regulation, and transport.

Finally, we examined the frequency of phenotypic alterations resulting from the knockdown of all CDG genes across the observed phenotypes. In this analysis, we compared the frequencies of phenotypic changes following the knockdown of the 23 CDG genes present within the network (“CDGs IN-Network”) to those resulting from the knockdown of the 14 CDG genes not part of the network (“CDGs OUT-Network”). It was observed that silencing in-network CDG genes significantly increased the likelihood of inhibiting biofilm-associated phenotypes, including biovolume, biomass, maximum height, mean thickness, and swarming motility (Fig.4D, top panel, Table S6). Conversely, the knockdown of out-network CDG genes tended to increase biofilm volume and mass, enhance swarming motility, and reduce the secretion of amyloid components. This divergent phenotypic behavior indicates that in-network CDGs are more likely to exert a positive effect on biofilm formation than those outside the network. Considering that most in-network CDG proteins are associated with DipA (PFLU0458), we compared the frequencies of phenotypic changes following the knockout of genes within the DipA interaction module to those of the out-of-network CDG genes (Fig. 4D middle panel, Table S6). The silencing of DipA-interacting partners was statistically more likely to inhibit several biofilm-associated phenotypes and swarming, underscoring the critical roles of DipA and its CDG partners in promoting biofilm formation. Finally, this trend persisted when the entire set of 56 genes from the PPI network was compared with the out-network set of 14 genes, indicating that CDG proteins and their network partners are significantly enriched in functions that promote biofilm formation (Fig. 4D bottom panel, Table S6).

#### 3.3.3. Global analysis of CDG proteins in SBW25 revealed phenotypic clusters related to biofilm and motility

Analysis of the phenotypic changes resulting from the knockdown of all in-network CDG proteins (Table S6) revealed a wide spectrum of biofilm and swarming phenotypes (Fig. S5). In our screens, we found that biofilm (BV) and swarming motility (SW) phenotypes were not necessarily inversely correlated, suggesting that they might not be antagonistic^78^ (Fig. S5A, B). Furthermore, CDGs exhibited other biofilm-related phenotypes (BV and SBP) that did not necessarily co-occur (Fig. S5C). Examination of biofilm volume showed that silencing the expression of 20 out of 37 CDGs resulted in increased biofilm volume, whereas 12 were impaired in biofilm formation. In general, biofilm formation or inhibition was not correlated with the composition of the CDG domains (Fig.S5A). Notably, four CDGs demonstrated a strong negative impact on biofilm production when downregulated (Log2Fold from −3 to −4.6). Among these, DcgP (PFLU0085), GcbA (PFLU0621) and GcbC (PFLU956) have previously characterized DGC activities (Table S1), and the PilZ protein Alg44 (PFLU0988) is involved in the production of alginate, thereby supporting their prominent role in biofilm formation. Knockdown of PDE RimA (PFLU0263) also significantly reduced biofilm formation, suggesting that the downregulation of enzymes with antagonistic activities on c-di-GMP could result in a similar biofilm reduction phenotype. BifA (PFLU4858) and GcbB (PFLU4600) were mainly involved in swarming activity (Fig. S5B). Knockdown of *bifA* also resulted in the greatest increase in submerged biofilms at the pegs, in sharp contrast to *dipA*, which exhibited the most significant decrease in SBP upon knockdown (Fig.S5C). Our observations align with previous studies on other *Pseudomonas* species, which also highlighted the hyperadhesive phenotype of *bifA* mutant cells^43,79^. However, the low adhesive capability of cells with *dipA* knockdown contrasts with the previously characterized role of *dipA* in biofilm dispersal and cell attachment in *P. aeruginosa* ^42,80^.

Hierarchical clustering of standardized Log2Fold changes across all phenotypes revealed distinct groups of CDG gene knockdowns with comparable phenotypic signatures (Fig. S6). All CDGs exhibited negative phenotypes for one or more biofilm and motility traits. DcgP (PFLU0085), GcbA (PFLU0621), GcbC (PFLU956), MucR (PFLU2753), and the PilZ protein Alg44 (PFLU0988) formed a clear cluster with a strong negative impact on biofilm volume and mass upon silencing, indicating that they play a role in biofilm formation. A group of 13 CDGs exhibited reduced swarming capacity and increased submerged biofilm upon knockdown, suggesting their involvement in the transition from motile to sessile states, such as during biofilm dispersal. This cluster includes RapA (PFLU2031), BifA (PFLU4858), MorA (PFLU5329), GcbB (PFLU4600) and other uncharacterized CDGs. This behavior is consistent with the phenotypes of BifA in other *Pseudomonas* species^43,79^. Finally, the third cluster comprised six CDGs, including AdrA (PFLU3650) and AwsR (PFLU5210), for which knockdown resulted in defects in amyloid secretion. This suggests their potential role in regulating the secretion of extracellular matrix (ECM) components.

Principal component analysis (PCA) was conducted on various parameters associated with biofilm pellicle, swarming, and extracellular amyloid production (Fig. 5A). Clustering analysis identified three principal components that accounted for 86% of the variance (Table S7-1). The first two components explained 46.5% and 22.4% of the variance, respectively. The primary factors contributing to PC1 were related to biofilm mass (thickness and height), whereas roughness and amyloid polysaccharides production were mainly associated with PC2 (Table S7-1). The third component was characterized by swarming behavior. Hierarchical clustering analyses delineated three distinct clusters of phenotypic variation. Clusters 1 and 2 were largely separated from Cluster 3 by the first principal component, whereas the second component separated Clusters 1 and 3 from Cluster 2 (Fig. 5A, Table S7-1). The CDG proteins were distributed in the clusters irrespective to their GGDEF/EAL/PilZ domain structural organization (Fig.5A). Cluster 1 included five interaction partners inversely correlated with DipA (PFLU0458), namely DgcP (PFLU0085), GcbA (PFLU0621), GcbC (PFLU0956), WspR (PFLU1225), and MucR (PFLU2743), in addition to the PilZ-domain protein Alg44 (PFLU0988) and the phosphodiesterase RimA. Cells depleted for these CDGs exhibited a strong defect in biofilm production (Fig. S5B). Cluster 2 comprised 13 members, including DipA and six of its interaction partners, such as RbdA (PFLU4308), RapA (PFLU2031), and MorA (PFLU5329). It was enriched in CDGs genes not present in the network (nine of 14 out-of network targets). Cluster 3 consisted of 17 CDG genes, whose depletion increased biofilm production while decreasing swarming motility (Fig. S5B). This cluster included four CDG baits: BifA (PFLU4858), PFLU5127, PFLU5210, and PFLU1114.

**Figure 5:**
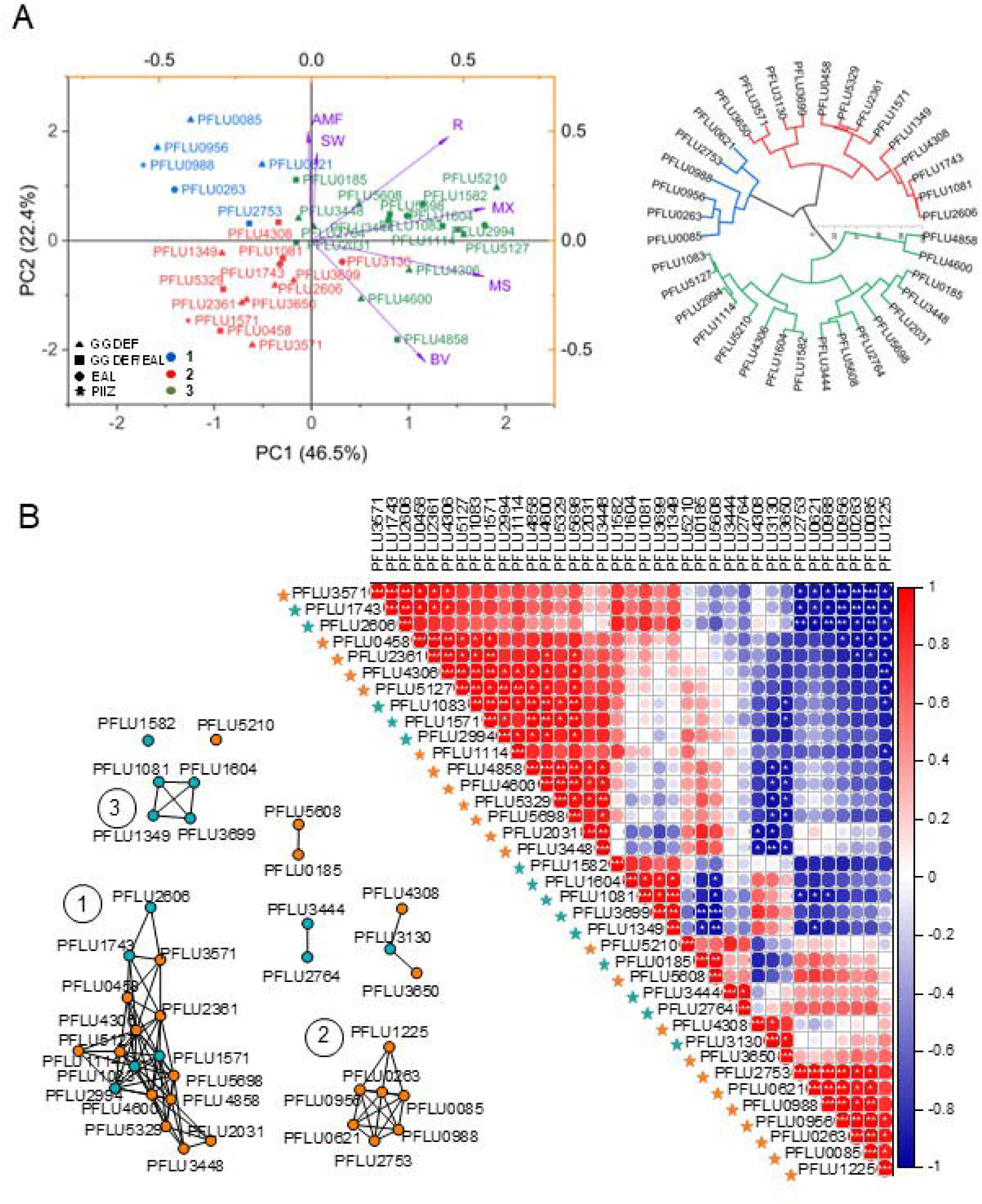
Phenotypic correlation analysis between all CDG genes. A- Principal component analysis plot with K-means clustering of biofilm pellicle and swarming motility phenotypes. Note: the less robust SBP phenotype was excluded from this analysis; WspR (PFLU1225) is here missing due to the lack of AMF data). Clusters are indicated by colors (1= green, 2= blue and 3= red). Purple arrows indicate the eigenvectors associated with the biofilm and swarming phenotypes. B- Pearson correlation analysis between Log_2_ fold change of the biofilm and motility phenotypes (BV, MX, MS, R, SW, and AMF). Positive and negative correlations are indicated by red and blue shades, respectively. The genes were ordered to group the most correlated. “In-network” vs “out-network CDG genes are indicated by orange and blue asterisks, respectively. The vertical bar represents the color legend of the correlation coefficients. The size of the dot also reflects the value of the correlation coefficient. White asterisks indicate the statistical significance levels (* p ≤ 0.05, ** p ≤ 0.01, *** p ≤ 0.001). Left: edge-weighted network of significant correlations. Each node represents a CDG gene from the PPI network (orange) or outside the network (blue).

Pearson correlation coefficients (PCC) between gene pairs also revealed groups of phenotypically correlated CDGs (Fig. 5B, Table S7-2). Note that the SBP phenotype was excluded from the analysis due to the reduced robustness of the protocol when the material attached to the pegs was insufficient (see Materials and Methods). The main group (group 1) comprised 17 genes, including DipA (PFLU0458), and 10 of its interaction partners. This group was enriched with IN-network CDGs (12/17). Group 2 contained seven members, all belonging to the network, which strictly corresponded to the members of PCA cluster1. Similar to the PCA, the members of group 2 were significantly anti-correlated with a subset of genes in group 1. The third group consists of four inter-correlated OUT-network CDGs, which are the putative PDEs, EAL-containing proteins PFLU1604, PFLU1349, PFLU1081, and PFLU3699, the latter two containing degenerate GGDEF signatures. This group 3 was anti-correlated with DGC and GGDEF-containing proteins, PFLU5608 and DgcH (PFLU0185).

In summary, the global overview of phenotypic patterns revealed a non-random distribution with marked asymmetry for key traits between the CDGs IN- and OUT-networks (Fig. 4D). CDGs can be categorized into three distinct clusters based on their phenotypic correlations (Fig. 5). The network was enriched in CDGs, for which silencing led to the suppression of biofilm and/or motility traits, indicating that they positively regulate biofilm and/or motility phenotypes. Conversely, the knockdown of OUT-network CDGs was more closely linked to phenotypes resulting from the activation of biofilm traits, indicating a negative regulation of biofilm formation. The intermediate group exhibited mixed profiles that potentially corresponded to modulators or signal integrators. These results emphasize the complex mechanisms that regulate the development of various biofilm types (i.e., air-liquid pellicle or solid-liquid submerged) and the composition of the extracellular matrix.

#### 3.3.4. A detailed examination of the functional relationships between key CDG proteins and their interaction partners

First, we focused on the DipA (PFLU0458) interaction module because of its pivotal position within the c-di-GMP signaling network. Pearson correlation coefficients (PCC) between gene pairs within DipA subnetwork revealed groups of phenotypically correlated genes (Table S7-3). This analysis revealed 3 main groups of CDG proteins with phenotypes positively and negatively correlated with DipA (Fig. 6A). The first group, composed of DipA and the 4 uncharacterized CDGs PFLU2361, PFLU4306, PFLU5127 and PFLU3571 exhibited phenotypes strongly correlated with each other and significantly negatively correlated with the 3 characterized CDGs PFLU0956 (GcbC), PFLU0085 (DgcP) and PFLU1225 (WspR). A representation of phenotype changes upon knockdown of genes from the DipA module showed genes in this first group exhibited increased biofilm volume and reduced swarming mobility (Fig. S7A). This observation suggests that DipA may be part of a group of CDG proteins that negatively regulate biofilm formation and positively regulate swarming motility. The second phenotypic group consisted of seven highly inter-correlated partners that were not significantly correlated with DipA. This group included five CDGs, namely PFLU4600 (GcbB), PFLU5329 (MorA), PFLU5698, PFLU2031 (RapA), and PFLU3448, as well as two non-CDG genes PFLU4971 and PFLU4159. The third phenotypic group comprised three characterized CDGs, namely PFLU1225 (WspR), PFLU0085 (DgcP), and PFLU0956 (GcbC), which strongly impacted biofilm formation upon silencing and exhibited phenotypes significantly negatively correlated with those of the DipA cluster (Fig.6A, Table S7-3). PCA identified three principal components (PCs) that accounted for 95% of the variance (Fig. 6B, Table S7-4). The first two components explained 52% and 31% of the variance in this study, respectively. The primary factors contributing to PC1 were biofilm volume, height, and thickness, which are directly associated with biofilm mass. PC2 differentiated gene phenotypes primarily based on biofilm roughness and swarming motility. The main contributing factor for PC3 (12%) was swarming. K-means clustering defined three clusters that largely overlapped with the three PCC groups identified in Fig. 6A. The first cluster (blue in Fig. 6B) contained five members whose phenotypes showed the strongest inverse correlation with DipA. The second cluster (red in Fig. 6B) included five members, among them DipA, PFLU2361, and PFLU3571, with phenotypes showing strong positive correlations, as well as PFLU5329, which belongs to the second PCC group. The third cluster (black in Fig. 6B) consisted of nine members, including six of the seven members of the second PCC group. The combined PCC and PCA analyses highlighted the existence of both positive and negative functional relationships between DipA and its interacting partners in biofilm formation and swarming motility.

**Figure 6:**
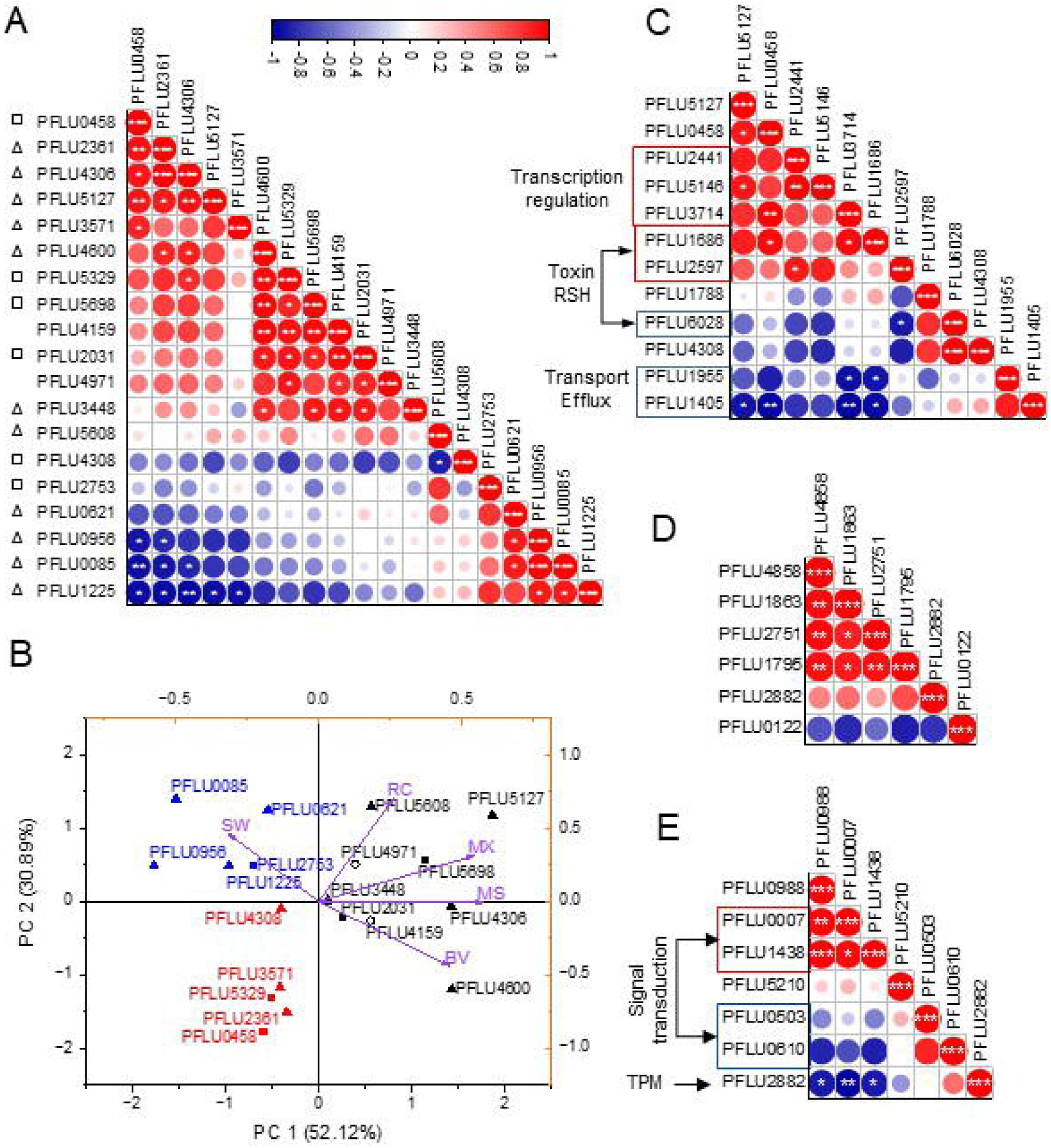
Phenotypic correlation analysis of gene pairs. A- dipA interaction module. Pearson correlation matrix for Log2FC in biofilm and motility phenotypes (BV, MX, MS, R, SW, and AMF). Positive and negative correlations are indicated in shades of red and blue respectively. Genes were ordered from the highest to lowest correlation with dipA. The color intensity and size of the circles are proportional to the correlation coefficients. The vertical bar shows the color legend of the correlation coefficients. Significantly correlated positive and negative clusters are indicated by red and blue shades, respectively. White asterisks indicate the levels of statistical significance (* p ≤ 0.05, ** p ≤ 0.01, *** p ≤ 0.001). Domain composition is indicated (GGDEF, open triangle; GGDEG/EAL, open square). B- Principal component analysis plot with K-means clustering of genes based on biofilm and motility phenotypes. Clusters are indicated by colors (1 red, 2 blue and 3 black). The eigenvectors associated with the biofilm and swarming phenotypes are shown in purple. CDG proteins containing the GGEEF domain (filled triangle) or dual GGDEF/EAL domain (filled square). Open circles are non-CDG partners. C- PFLU5127 interaction module. Pearson correlation matrix for Log2FC in biofilm and motility phenotypes (BV, MX, MS, R, SBP, and AMF). D- BifA (PFLU4858) interaction module. Pearson correlation matrix for Log_2_ fold change in biofilm and motility phenotypes (BV, MX, MS, R, and SW). E- Alg44 (PFLU0988) interaction module. Pearson correlation matrix for Log_2_ fold change in biofilm and motility phenotypes (BV, MX, MS, R, and SW). Partners with similar functional annotations are framed in red (correlated) or blue (anti-correlated).

PFLU5127 and 11 of its interacting partners for which phenotypes were measured, were analyzed similarly (Fig.6C, Fig. S7B, Table S7-5). Three transcriptional regulators PFLU3714, PFLU5146 and PFLU2441, together with DipA, positively correlated with the PFLU5127 phenotypes. Conversely, PFLU5127 displayed an inverse correlation with PFLU1405 and PFLU1955, which encode a TonB-like receptor and a porin involved in the uptake and transport of components and nutrients, respectively. These findings underscore the potential role of c-di-GMP in modulating transcription and substrate transport during the formation of biofilm. Phenotypic correlations between PFLU5127 and three out of four targeted interacting partners of the RHS-superfamily of proteins were not significant for the phenotypes examined. In addition, examination of phenotype changes further indicated an inverse relationship between the regulation of biofilm and swarming phenotypes by CDGs in this module (Fig. S7B).

Regarding the phosphodiesterase BifA (PFLU4858) and its 5 high confidence interacting partners for which phenotypes were measured (Fig.6D, Table S7-6), our analysis revealed a significant positive cluster comprising the transcriptional regulator PFLU1863, efflux pump component PFLU2751, phage tail-like bacteriocin PFLU2882, and exported protein PFLU1795. In contrast, an inverse correlation was observed with the transcriptional regulator PFLU0122. Depletion of *bifA* resulted in a marked increase in biofilm formation and a defect in swarming motility (Fig.S7D), consistent with previous studies ^43,77^. In addition, we found that *bifA* knockdown led to the highest increase in submerged biofilms at pegs (Fig. S5C, SBP), while producing significantly fewer exopolymeric amyloids (Table S5). This suggests an enhancement in cell adhesion, in keeping with the hyper-adherent phenotype observed in other *Pseudomonas* strains^79^.

Finally, the PilZ protein Alg44 (PFLU0988) was significantly positively correlated with the histidine kinase sensor protein PFLU0007 and the methyl chemotaxis protein PFLU1438 (Fig.6E, Fig. S7D, Table S7-7), and was negatively correlated with phage tail-like bacteriocin PFLU2882. Taken together, our results strongly support a functional relationship between CDGs and their protein partners as well as a role of c-di-GMP in the coordination of several biological pathways including regulation of gene expression, signaling, interbacterial competition, transport and adhesion to surfaces.

### 3.4. Exploration of the Multifaceted Functions of c-di-GMP Signaling

#### 3.4.1. A rapid screening of the subcellular localization of c-di-GMP binding proteins

Previous studies have demonstrated that a discrete pattern of subcellular localization can be advantageously used to establish the functional relevance of protein-protein interactions^81^. Thus, we set out to screen for CDG proteins exhibiting discrete subcellular localizations, focusing on our most striking observations: (i) the DipA subnetwork connects 17 CDGs proteins, 11 of which contain a GGDEF domain, with 8/11 characterized as functional diguanylate cyclases in *Pseudomonas* (Table S1), and 6 (including DipA) containing a dual GGDEF/EAL domain, with 5/6 characterized as PDE or dual PDE and DGC activities in *Pseudomonas* (Fig. 2A, Table S1); and (ii) the connection of CDGs with numerous cell processes, including cell division and DNA repair. The C-terminal ends of 16 CDG proteins were fused to the mNeonGreen fluorescent protein, and the fusion proteins were expressed under the control of the rhamnose-inducible P_BAD_ promoter, as described in Materials and Methods., Discrete patterns of subcellular localization were observed for 14 different fusion proteins expressed during exponential growth (Fig.S8). PFLU1114, along with DipA and four of its partners, formed a single focus localized at one cell pole, which could reflect a biologically relevant localization or a tendency of the fusion proteins to accumulate at the pole. In contrast, PFLU5127 exhibited a bipolar localization, and PFLU5608 displayed a single focus at one pole in short cells and appeared to form a ring-like structure in the middle of longer cells, suggesting a septal localization (Fig. S8). Notably, RimA exhibited a single focus per cell at various positions, suggesting potentially dynamic localization. Finally, among the proteins with membrane-associated localization, Alg44, AdrA, PFLU0007, and PFLU0610 localized as diffuse crescent-like shapes covering both poles of the cell, whereas PFLU0503 was more homogenously distributed in the cell membrane. This crescent-like subcellular localization at the cell pole is reminiscent of the localization of chemoreceptor proteins in other bacteria^82,83^. Altogether, our screening for discrete subcellular localization provided us with three candidate CDG proteins, which we subsequently investigated further to ascertain their biological function, as inferred from the network: PFLU5608, AdrA (PFLU3650), and RimA (PFLU0263).

#### 3.4.2. PFLU5608 acts during cell division

PFLU5608 comprises a multipass membrane domain at the N-terminus and a cytoplasmic GGDEF domain at the C-terminus, suggesting it is a membrane-anchored protein with putative diguanylate cyclase activity (Fig. 2A). The septal localization of PFLU5608 C-terminally tagged with mNeonGreen fluorescent protein (PFLU5608-FP), observed in our screen, suggests PFLU5608 could play a role during cell division. Detailed analysis confirmed that PFLU5608-FP exhibited two types of localizations: as a ring at the septum of dividing cells and at one or both cell poles in divided shorter single cells (Fig.7A and Fig. S9, Movie S1). This localization pattern is highly evocative of that observed for other membrane-anchored proteins of the bacterial divisome. Notable examples include ZipA and FtsA, which have been shown to bind to the main component of the septal ring, FtsZ, thereby stabilizing the Z-ring in *E. coli* and other Gram-negative bacteria ^84,85^. To assess whether PFLU5608 is involved in regulating cell division in SBW25, we examined the effect of its absence on cell morphology using fluorescence confocal microscopy (Fig.7B). Compared to the wild-type SBW25 strain, the ΔPFLU5608 mutant strain exhibited several morphological defects, producing smaller cells and fewer elongated cells after overnight growth in liquid culture. To further investigate the role of PFLU5608 during exponential growth, we examined the morphological phenotype of cells in which PFLU5608 expression was silenced by CRISPR interference. Using our Tet-ON inducible CRISPRi/dcas9 system, we expressed a guide RNA targeting the PFLU5608 gene (g5608) (Table S5). Elongated cells were observed three hours post-expression of the dCas9/g5608 complex, indicating a defect in cytokinesis (Fig. 7B). Statistical analysis of cell size indicated moderate but significant cell elongation following PFLU5608 depletion (Fig.7C). The findings demonstrate that CDG PFLU5608 activity is important for a normal cell division during exponential growth of SBW25, although it is not essential. This establishes a functional link between cyclic di-GMP signaling and the regulation of cytokinesis in SBW25.

**Figure 7.**
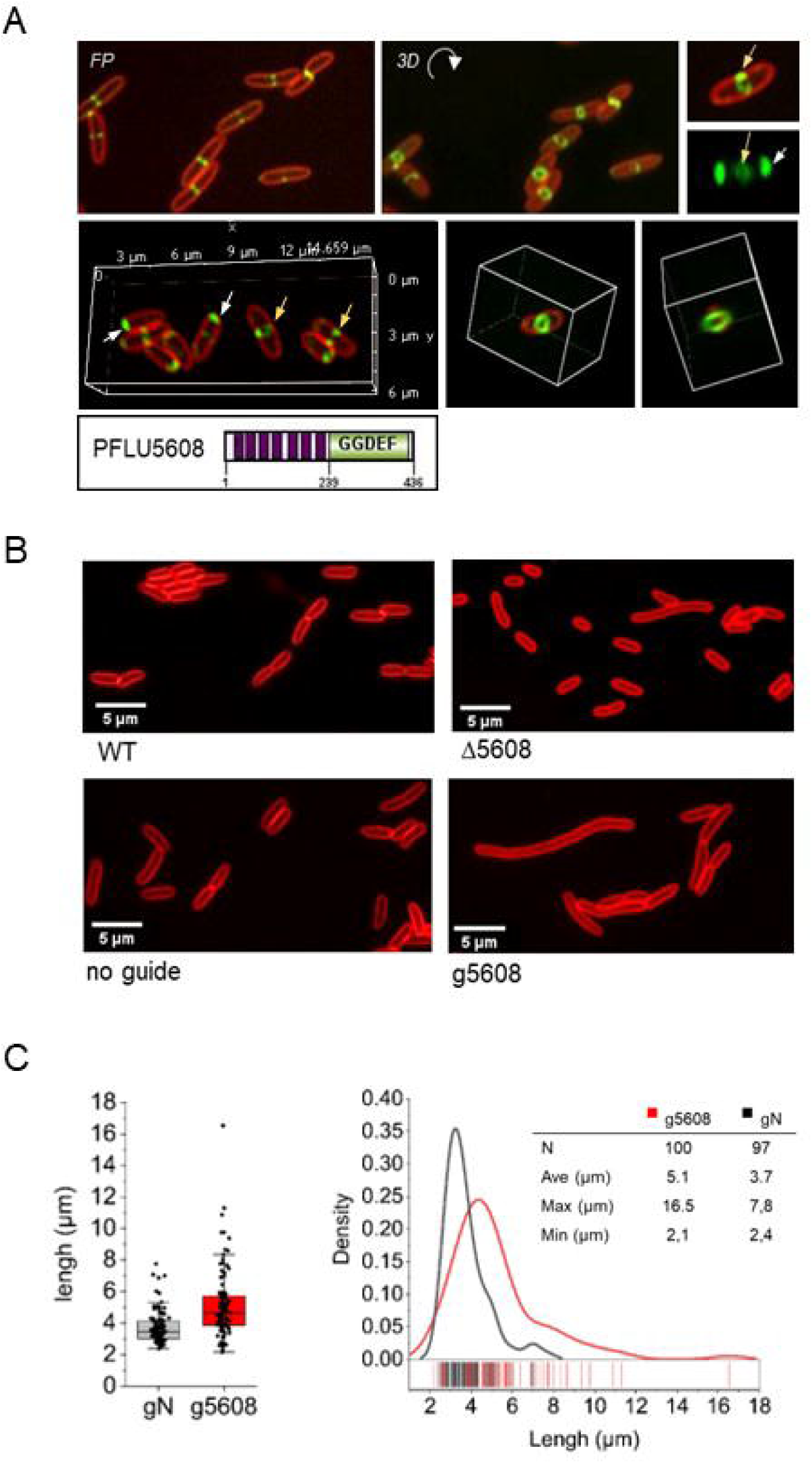
PFL5608 plays a role in cell division. A- Intracellular localization of the PFLU5608-mNeonGreen protein. 2D focal plane (FP) and 3D reconstructions showing PFLU5608 localizing at mid cell and forming a ring (yellow arrows) and at the cell poles (white arrows). Cell membranes were stained with the lipophilic dye FM 4-64 (red). B- Effect of PFLU5608 depletion of on cell length. A comparison was made between knockdown strain (g5608) and knockout strain (Δ5608). Cell membranes were stained with FM 4-64 (red). C- Cell counts and cell length measurements in knockdown strains expressing a guide targeting PFLU5608 (g5608) or a non-targeting guide sequence (gNT) were performed using Fiji with the MicrobeJ plug-in.

#### 3.4.3. AdrA acts in DNA repair

In our PPI network, AdrA (PFLU3650) interacts with PFLU5498 and PFLU3600, two UvrA homologs. UvrA is a component of the prokaryotic nucleotide excision repair (NER) complex UvrABC, which is responsible for detecting and excising lesions in damaged DNA, including DNA crosslink adducts^86^. A detailed examination of their binding domains revealed that the cytosolic GGDEF domain of AdrA, used as bait in our Y2H screen, interacted with the ABC-type ATP-binding domain of both UvrA homologs (Fig. 8A). We investigated the potential involvement of AdrA and its partners in the NER pathway of SWB25 *in vivo* by downregulating its expression along with that of each *uvrA*-like genes using CRISPRi, and assessed the bacterial survival after treatment with mitomycin C (MMC), a DNA-damaging agent that creates crosslinked adducts known to be repaired by NER (Fig. 8B, Table S5). Following exposure to 0.5 µg/ml of MMC, a decline in viability was observed in wild-type cells, attributable to an excess of DNA damage. This was observed in both the control strain that did not express guide RNA (2.8% survival rate) and in cells that expressed a guide that targets PFLU0085, an unrelated GGDEF protein used here as a negative control for NER (4.6% survival rate). However, cell survival was significantly reduced in the presence of MMC upon silencing of PFLU5498 and PFLU3600, with a 14-fold (0.33%) and 10-fold (0, 48%) decrease compared to the control, respectively (Fig. 8B). These findings corroborate the role of both UvrA homologs in NER, with PFLU5498 and PFLU3600 exhibiting non-redundant functions. Notably, silencing of PFLU3650 also resulted in a similar 14-fold reduction in MMC survival (0.32%). These findings are consistent with AdrA contributing to NER through its interaction with an UvrA-like protein, leading to the formation of a protein complex which could either modulate NER activity or direct it to specific sites within the cell.

**Figure 8.**
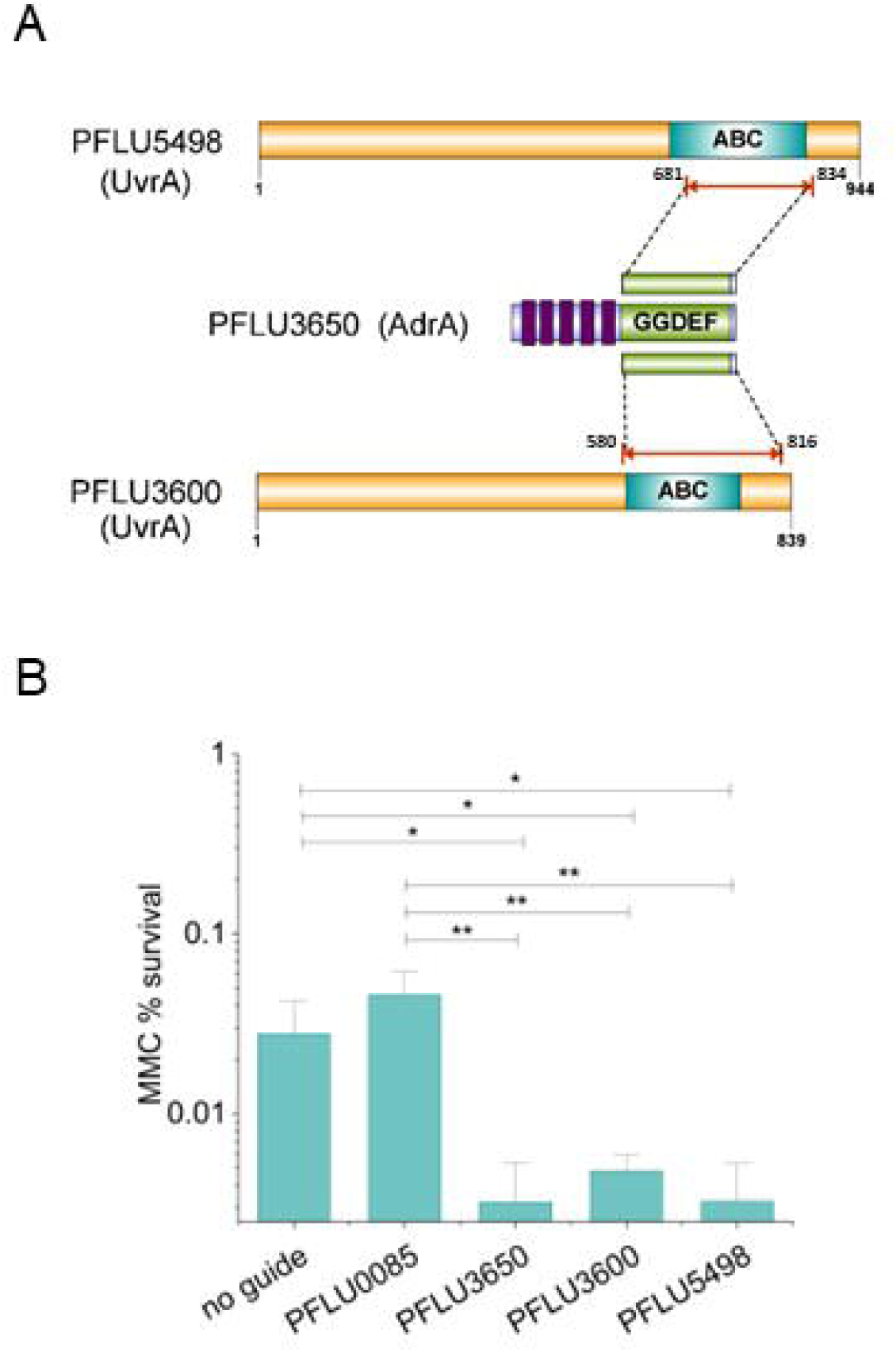
Functional validation of the role of PFLU3650 in DNA repair. A- Interacting domains with UvrA-like protein partners PFLU3600 and PFLU5498. B- Survival following exposure to mitomycin C (0.5 µg/ml) after CRISPRi-mediated knockdown of PFLU3650 or its interacting partners. The control was an isogenic SBW25 strain that did not express a guide RNA. A second control comprised a strain expressing a guide RNA targeting the cytosolic DGC DgcP (PFLU0085).

#### 3.4.4. Highly dynamic localization of RimA depends on a functional c-di-GMP binding site

The distinctive subcellular localization pattern of RimA attracted our attention. We determined that RimA-FP formed a single focal point per cell, which dynamically transitioned from one pole to the other in approximately 20 seconds. (Fig. 9A, Movie S2). Particle tracking was performed on 27 foci, revealing a displacement speed ranging from 0.2 to 1.12 µm/s, with an average speed of 0.26 µm/s (Fig. 9B). To investigate whether the phosphodiesterase (PDE) activity of RimA was involved in its dynamic movement, we constructed a RimA E47A mutant that specifically inactivated the EAL motif responsible for c-di-GMP hydrolysis. We observed that the RimAE47A-FP mutant was uniformly distributed in the cytoplasm and did not form dynamic foci in cells (Fig. 9B). In our assay, the FP-tagged mutated copy of RimA was expressed from a plasmid in cells containing a wild-type chromosomal copy of *rimA*, raising the possibility that the RimA-FP focus also contained wild-type RimA. However, the dynamic behavior of RimA-FP is abolished by a mutation inactivating the EAL domain of RimA, suggesting that RimA-FP dynamics may require the PDE activity.

**Figure 9:**
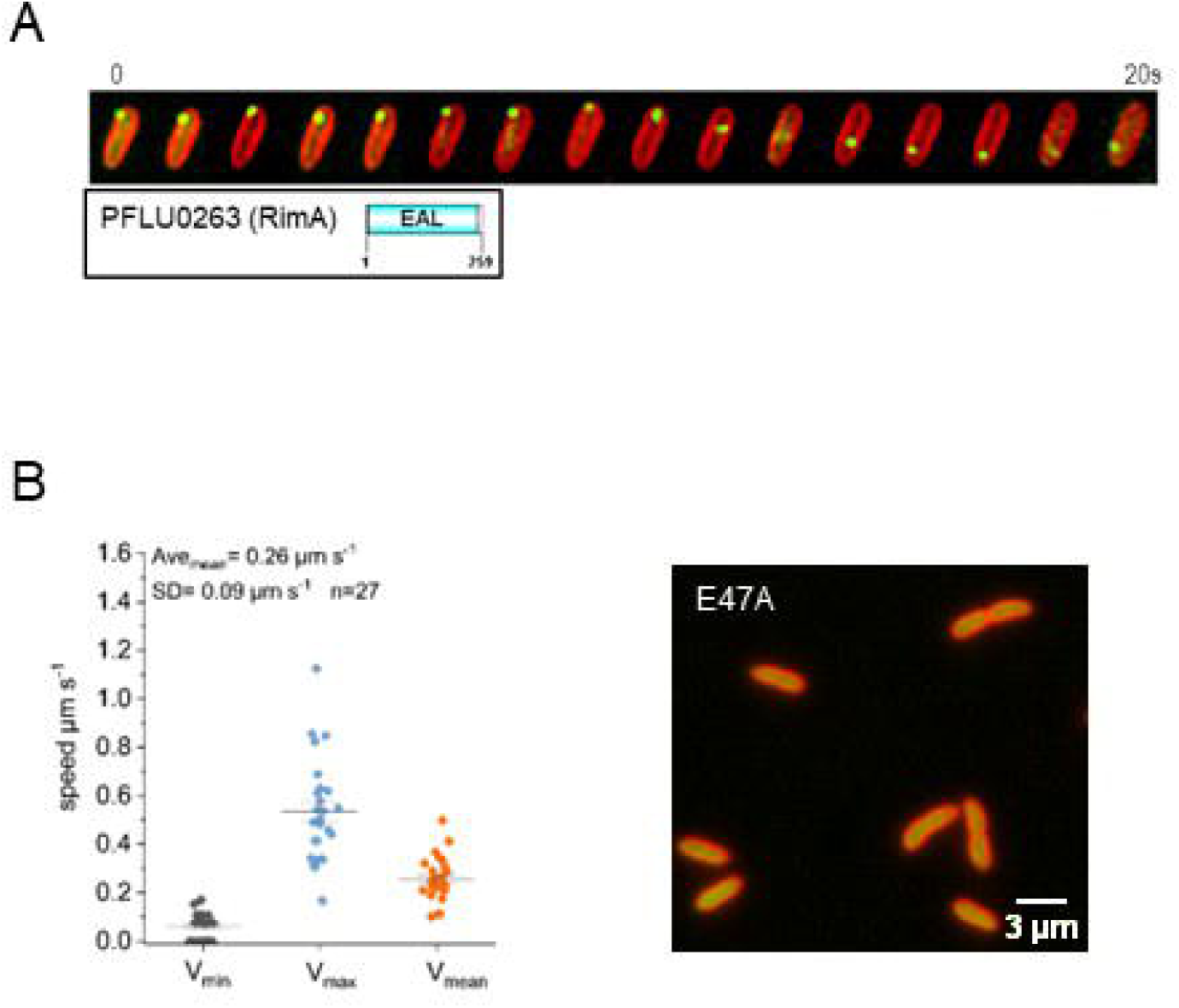
Analysis of RimA subcellular dynamics. A- Cells expressing a RimA-FP fusion protein from a plasmid exhibit a single dynamic fluorescent focus that scans the entire cell along its long axis. B-Average speed (µm/s) of RimA-GFP foci in individual cells (n=27, left). RimA dynamic behavior depends on the integrity of the EAL catalytic site (right). The RimA-FP protein containing an E47A substitution (EAL ◊ AAL) was homogenously dispersed in the cell and unable to form discrete foci.

## 4. Discussion

The c-di-GMP signaling pathway is integral to numerous bacterial cellular processes. In *Pseudomonas*, as in many other bacterial species, it is pivotal in the transition from a motile lifestyle to biofilm formation^12,13^. Various studies have also underscored the role of c-di-GMP in regulating other metabolic and cellular processes ^14,15,87–91^. However, much remains to be discovered about the metabolic pathways involved in this intricate developmental process, particularly their roles in driving biofilm formation. In this study, we adopted an integrated, multifaceted approach that combined genome-wide yeast two-hybrid screenings to identify interaction partners, with the examination of multiple phenotypic traits associated with biofilm formation and swarming motility after CRISPRi-mediated silencing of genes. Our objective was to deepen our understanding of the intricate c-di-GMP signaling pathway and the functions of CDG-proteins and their interacting partners. This study provides a foundation for developing hypotheses regarding the functional roles of CDGs in modulating different cellular pathways that may be involved in biofilm formation, such as those occurring in the rhizosphere.

We built a high confidence protein-protein interaction network centered on 10 CDGs that robustly connects the c-di-GMP signaling regulatory pathway to a variety of other cellular processes potentially implicated in biofilm development. In addition to c-di-GMP signaling pathway, the network identified novel interactions with two-component systems that mediate signaling through the phosphorylation of specific proteins. This finding underscores a significant interplay between c-di-GMP and other signaling pathways, as previously observed in other bacterial species ^92–94^. The network also extended to the transport and secretion systems of various compounds, the regulation of transcription, cell-cell adhesion and contact inhibition, as well as pathways related to cell division and DNA repair. A subset of CDG baits exhibited a high degree of connectivity with protein associated with specific functional categories. For instance, the analysis of subnetworks revealed significant associations between DipA and the c-di-GMP signaling pathway, Alg44 and two-component signal transduction systems (TCS), BifA and nutrient transport, PFLU5127 and toxin-antitoxin (TA) systems as well as between AdrA and DNA repair pathways. This finding suggests that these CDGs may have some degree of functional specialization in their ability to modulate various cellular processes.

In the network, BifA (PFLU4858) is primarily linked to functions such as multidrug efflux pumps, transport mechanisms and transcriptional regulators. Silencing *bif*A resulted in significant impairment of swarming and in a large increase of submerged biofilm development. This observation aligns with earlier findings in *P. aeruginosa*, where BifA inversely controls biofilm formation and swarming motility ^43^. Analysis of the phenotypic enrichment of CDGs and their interacting partners revealed their differential involvement in biofilm formation (Fig. 4). More specifically, silencing of RimA (PFLU0263) and DipA (PFLU0458) and their interacting partners was significantly associated with negative biofilm phenotypes, indicating that these proteins act to promote biofilm formation. In contrast, the silencing of RbdA (PFLU4308), BifA (PFLU4858), AswR (PFLU5210), PFLU5127, and their interaction partners were associated with a significant increase in biofilm phenotypes. This indicates that these CDG proteins act to repress biofilm formation and likely operate in the same regulatory pathway as a significant proportion of their interacting partners.

The examination of pairwise correlations between phenotypes of CDGs and their partners revealed both positive and negative associations (Fig. 6). A compelling example is the analysis of the phenotypic pattern of DipA (PFLU0458) knockdown, which showed a strong correlation with a cluster of four uncharacterized CDG partners, namely PFLU4306, PFLU2361, PFLU5127, and PFLU3571. In contrast, knockdown of three characterized DGCs, WspR (PFLU1225), DgcP (PFLU0085), and GcbC (PFLU0956), resulted in a significant anti-correlation with DipA. A similar observation was made for PFLU5127, which displayed significant positive correlations with two toxin-antitoxin delivery proteins and three transcriptional regulators, alongside a significant negative association with proteins involved in transport and efflux. Overall, these results suggest that CDG proteins can modulate biofilm and motility phenotypes, at least in part, through interactions with proteins that have either a positive or a negative impact on these phenotypes.

We observed also that the knockdown of IN-network CDGs was statistically more likely to result in decreased biofilm and swarming motility phenotypes, whereas the knockdown of OUT-network CDGs was statistically more likely to result in increased phenotypes (Fig. 4C). It is apparent that DipA and its multiple CDG interacting partners have a substantial impact on this observation. However, the enrichment was still significant when considering all 70 proteins in the network (i.e. CDGs and partners) that were phenotyped. Another relevant factor is our initial selection of 10 CDG baits exhibiting functional roles at various stages of biofilm formation in root-associated *Pseudomonas* species. This combination of factors may explain the network’s enrichment in proteins that contribute positively to biofilm formation compared to those outside the network. Our phenotypic screens also revealed that the complement of CDG genes formed three highly correlated clusters (Fig. 5B), suggesting the potential formation of functional CDG modules with members acting similarly in biofilm formation.

### DipA as a c-di-GMP central hub

In our PPI network, it is remarkable that 16 out of 18 DipA (PFLU0458) partners are also CDG proteins. Furthermore, the specific interaction domains identified in Y2H genomic screens corresponded to the GGDEF domains for all these proteins, indicating the potential formation of heterodimeric complexes with the corresponding domain of DipA, which contains a degenerated ASNEF motif. These observations suggest that DipA functions as a central hub for c-di-GMP interactions. The presence of a single interaction GGDEF domain common among DipA partners implies that DipA could be classified as a “singlish hub” (previously termed “date hub”), forming transient protein complexes with its partners at different times within the cell, as opposed to a “multi-interface hub” that interacts with multiple partners simultaneously through distinct interaction domains^95^. The identification of such a highly connected CDG hub aligns with other interaction studies conducted on *P. fluorescens* Pf0-1 and *E. coli*, which have demonstrated the high propensity of c-di-GMP-binding proteins to interact with each other^12,30^. A systematic analysis of all binary interactions between all DGCs and PDEs in *E. coli* identified a core of five hyperconnected CDGs, which are organized as “supermodules”, highlighting the formation of homologous and heterologous interactions between their GGDEF domains^12^. Another study revealed numerous GGDEF/EAL dual-domain and GGDEF single-domain proteins that formed a dense network of binary interactions between the 42 c-di-GMP binding proteins present in *P. fluorescens* Pf0-1^30^. However, in this comprehensive study, Pf0-1 DipA (pfl01_0460) did not interact with other CDGs, whereas the LapD protein physically contacted 12 GGDEF-containing putative DGCs. The LapADG signaling system, involved in cell adhesion in Pf0-1 and other *P. fluorescens* strains, is absent in SBW25. The present study revealed the high degree of promiscuity exhibited by SBW25 DipA towards numerous GGDEF-containing proteins. This finding, obtained through a genome wide Y2H screening approach, underscores the functional role of DipA as a c-di-GMP center. The parallel between the CDG hub in *P. fluorescens* SBW25 and other CDG hubs in *P. fluorescens* Pf0-1 suggests that c-di-GMP regulatory circuits consistently rely on the formation of a complex hub, which is not contingent on conserved proteins and interactions. Instead, the essential functions fulfilled by CDG hubs can be mediated by different proteins in various *Pseudomonas* strains, indicating a high degree of evolutionary adaptability of these proteins.

*P. aeruginosa* DipA is a dual GGDEF/EAL domain phosphodiesterase, involved in biofilm dispersion^42^. The absence of DipA is marked by reduced swarming motility and decreased exopolymeric matrix production. In SBW25, *dipA* knockdown resulted in a denser and smoother biofilm and impaired swarming^77^. The PDE RapA (PFLU2361) and potential DGCs PFLU4306 and PFLU5127, which displayed phenotypes strongly correlated with those of DipA, may similarly participate in biofilm dispersal. In SBW25, WspR is a response regulator DGC extensively characterized for its role in contact-dependent responses to solid surfaces and cellulose production^53,96^. DgcP, GcbA and GcbC were identified as contributors to biofilm development in a various *Pseudomonas* species^40,97,98^. In our study, these four diguanylate cyclases formed a highly phenotypically correlated cluster, inversely correlated to DipA. Together, these observations suggest that SBW25 DipA may function as a regulator and/or coordinator of the activities of multiple DGCs, which exhibit divergent effects on biofilm and motility phenotypes. In this context, considering the tendency of DipA and many CDGs proteins to localize at the cell pole, either as a single focus or as a diffuse crescent, we hypothesize the existence of a CDG center physically located at the cell poles that could locally modulate the c-di-GMP pool in the cells. In *P. aeruginosa* PA14, the DipA/Pch protein is localized to the flagellated cell pole by the chemotaxis machinery, generating asymmetry in intracellular c-di-GMP levels during the cell cycle^99^. In SBW25, DipA was also found to interact with a methyl-accepting chemotaxis protein PFLU5971. Collectively, our observations provide further evidence supporting a role of DipA in the regulation of local c-di-GMP pool size.

### Crosstalk between c-di-GMP signaling and other cell processes

Crosstalk between c-di-GMP and other cellular pathways is necessary for integrating diverse internal and external stimuli to adapt cellular responses to environmental conditions. In the network, the two-component sensor kinase PFLU0007 was found to be connected to four CDGs: Alg44 (PFLU0988), RbdA (PFLU4308), BifA (PFLU4858), and AswR (PFLU 5210). Knockdown of PFLU0007 severely impaired biofilm formation, similar to Alg44, highlighting a complex interplay between signaling via two-component systems and the c-di-GMP signaling pathway in biofilm development. However, PFLU0007 did not exhibit biofilm-related phenotypes correlated with the three other CDG-interacting partners, providing no information on the potential crosstalk with other pathways involving RbdA, BifA, and AswR. Another significant player is the adhesin activator PFLU0147, which exhibits a phenotypic profile highly congruent with that of PFLU0007. Both were connected by RbdA. In *P. aeruginosa*, RbdA is a membrane-associated PDE involved in biofilm dispersal, and a mutant inactivated for RbdA produces thicker biofilms^67^. In SBW25, RbdA silencing did not significantly change the phenotypic profile, suggesting that the function of RbdA is likely different in *P. fluorescens*.

Among the gene knockdowns that resulted in a complete loss of biofilm formation and swarming motility, the putative transcriptional regulator PFLU0122 was identified. PFLU0122 interacted with BifA (PFLU4858), and its phenotypic profile was inversely correlated with that of BifA. The importance of PFLU0122 in the production of exopolymeric amyloid fibers suggests that PFLU0122 might directly or indirectly regulate genes involved in the secretion of exopolymeric components. Similarly, PFLU0122 could regulate swarming motility by modulating genes associated with the flagellar machinery or the secretion of biosurfactants.

Notably, not all protein partners in the network exhibited correlated phenotypes with their CDG bait. This could be attributed to the potential of CDGs and their partners to perform a range of functions that extend beyond the scope of biofilm. From this perspective, we discovered that PFLU5608 assembles as a ring at mid-cell, and that knockout and knockdown of PFLU5608 resulted in modification of cell length (Fig. 7). This suggests potential crosstalk between c-di-GMP and the regulation of cytokinesis. The regulation of cell division by c-di-GMP has been already reported in other bacterial species. In *E. coli* and *Salmonella*, the DGC YfiN was also found to localize at mid-cell in an FtsZ-dependent manner, halting cell division by blocking cytokinesis in response to envelope stress^20^. However, in *P. fluorescens* SBW25, the YfiN/AwsR homolog (PFLU5210) did not show a discrete subcellular localization, suggesting that it may have other function. Supporting this observation is that within the network, PFLU5210 was highly connected to the two-component signaling systems. Future investigations will be required to determine the precise role of PFLU5608 in the division machinery.

Our PPi network also revealed several connections between the c-di-GMP pathway and DNA repair. We have confirmed that AdrA (PFLU3650) and its two UvrA-like interacting partners, PFLU3600 and PFLU5498, function within the NER pathway in SBW25 (Fig. 8). In *V. cholerae,* c-di-GMP is involved in cell tolerance to the mutagenic agent methyl methanesulfonate ^18^. In *B. subtilis*, the DNA damage response pathway is induced during biofilm formation, indicating that genes involved in DNA repair are being upregulated^100^. Regarding biofilm and swarming motility phenotypes, we found that in SBW25, knockdown of AdrA mainly affected the production of amyloids without significantly impacting swarming motility. In *S. enterica*, AdrA is involved in the regulation of exopolysaccharide production, whereas in the *P. fluorescens* F113, AdrA plays a role in flagellar motility^101,102^. Collectively, our results indicate that, in SBW25, the GGDEF-domain diguanylate cyclase AdrA serves a dual function, participating in both in c-di-GMP signaling and in DNA repair, suggesting that it may coordinate both processes during the biofilm life cycle. AdrA’s role in DNA repair is likely mediated by its physical interactions with two UvrA homologs. More broadly, AdrA exemplifies how highly conserved c-di-GMP signaling proteins can participate in diverse cellular processes across different bacterial species, indicating a significant degree of evolutionary flexibility in the wiring of c-di-GMP signaling pathways in each species.

Our study also uncovered the distinct dynamic behavior of the PDE RimA’s subcellular localization. Co-expressed from the *rimABK* operon, RimA and RimK, in conjunction with c-di-GMP, form a complex that regulates the level of glutamation of the ribosomal protein S6 (RspF), which in turn modulates ribosome function in response to environmental cues^103,104^. Within our network, RimA interacted mainly with proteins annotated in DNA metabolism and toxin secretion pathways. The potential involvement of RimA in these pathways remains to be confirmed. However, we showed that the dynamic pattern of RimA subcellular localization was disrupted upon alteration of its EAL motif, suggesting that this motion might be dependent on RimA’s phosphodiesterase activity. These characteristics closely resemble the dynamic scanning motion previously observed for *Bacillus subtilis* DisA, a diadenylate cyclase (DAC) that acts as a checkpoint to maintain genome integrity during spore development^105^. DisA localizes as a single focus moving rapidly across the cell long axis and pauses its movement upon induction of DNA damage^105^. DisA is composed of a DAC domain combined with a RuvA-like Holliday junction DNA-binding domain^106^. Further research revealed that the DNA recombination protein RecA displayed comparable behavior when *B. subtilis* cells entered sporulation^107^. The similarities observed in the dynamic patterns of DisA and RecA reflect DisA’s ability to interact with RecA nucleofilaments during DNA repair^108^. Based on this well-documented scanning motion by the c-di-AMP binding protein DisA, we propose that the dynamic PDE activity-dependent movement of SBW25 RimA could also reflect a scanning-like mechanism. However, the cell structure scanned by RimA remains an open question at this stage.

## 5. Concluding remarks

Our c-di-GMP-centered interactome identified many potential effectors associated with DGCs and PDEs, which are integral to functional classes relevant for biofilm formation, such as during the colonization of the rhizosphere by the plant-beneficial *P. fluorescens* SBW25. Although the biological roles of most CDGs and their interacting partners remain to be fully characterized, we confirmed the function of c-di-GMP-binding proteins in DNA repair and cell division pathways. Our protein-protein interaction network also highlighted a complex regulatory network with crosstalk between c-di-GMP and other signaling pathways, notably two-component systems. Most importantly, this study unveiled the existence of an interaction hub connecting the PDE DipA with at least 17 DGC proteins interacting through their GGDEF domains. We propose that DipA interaction facilitates the transient recruitment of DGCs to activate downstream effectors. In SBW25, this transient binding of multiple DGCs at DipA, potentially at the cell pole, may orchestrate the global output of the c-di-GMP signaling pathway. Future research will be required to fully understand the regulatory pathways controlled by c-di-GMP and their interconnexions with other signaling pathways in bacteria. However, our results strongly support the hub-based model of c-di-GMP signaling for localized control and signal integration ^60^, thereby facilitating rapid adaptation to environmental changes in a highly intricate signaling network.

## 6. Material and Methods

### 6.1. Bacterial strains and growth conditions

*P. fluorescens* SBW25 strain was obtained from G. Preston and P. Rainey (University of Oxford, ^52,109^). Plasmids were constructed and propagated in *E. coli* DH5α (Biolabs) as a cloning host prior to transformation into SBW25. Bacterial cultures were grown in LB medium or M9 medium supplemented with glucose 0.4% as the carbon source. When necessary, the selection for plasmid maintenance in *E. coli* and *P. fluorescens* was achieved by the addition of kanamycin (50 μg/ml) and/or gentamycin (10 μg/ml) to the growth media. Anhydrotetracycline (aTc, 100 ng/ml) was used as an inducer of the PtetA/TetR promoter/repressor system controlling the expression of *dcas9*. Cells were grown at 37 °C for *E. coli* and 28 °C for *P. fluorescens* under shaking conditions. All swarming and biofilm assays with *P. fluorescens* were performed in a humidity-controlled growth chamber at 25 °C and 70% humidity.

### 6.2. Yeast two-hybrid screenings

The Y2HB screens of *P. fluorescens* SBW25 genomic library were performed by Hybrigenics Services, S.A.S., Paris, France (https://www.hybrigenics-services.com) following the ULTImate Y2H^TM^ technology. Genes encoding the c-di-GMP-binding proteins PFLU4308 (RbdA), PFLU0458 (DifA), PFLU5127, PFLU5210 (AwsR), PFLU4858 (BifA), PFLU1114, PFLU0263 (RimA), PFLU0988 (Alg44), PFLU3650 (AdrA), and PFLU2031 were PCR-amplified from *P. fluorescens* SBW25 genomic DNA and cloned into pB27 and pB66 bait vectors allowing the translational fusion of the ORFs at the C-terminal end of the binding domains of LexA or Gal4, respectively. PFLU3650 was specifically designed to exclude N-terminal multi-spanning membrane domains. For library construction, 50 µg of genomic DNA from *P. fluorescens* SBW25 was fragmented by nebulization (Atomisor NL9M) to generate fragments of 1 kb size on average. DNA fragments were blunted by treatment with mung bean nuclease, T4 polymerase, and Klenow fragment prior to ligation onto the pP6 prey vector, allowing N-terminal fusions with the activation domain of Gal4^110^. The library was first transformed into *E. coli*, yielding 2×10^7^ clones containing independent fragments. Sequence analysis of 192 randomly chosen clones was performed to establish the library’s general characteristics. This primary library was then harvested and transformed into yeast to generate 5×10^6^ independent clones. The screening conditions were established for each bait by adjusting the selectivity of the HIS3 reporter with 3-aminotriazole (3AT) as described previously (Rain et al., 2001). All parameters of the performed screens are summarized in supplementary Table S2. All interactions were assigned a Predicted Biological Score (PBS) as described elsewhere^61,62^.

### 6.3. CRISPRi gRNA design and cloning

gRNAs were designed to target genes encoding c-di-GMP-binding proteins and their interaction partners as previously described ^57^. The 72 gRNA sequences (listed in Table S4) were synthesized as DNA gBlocks (IDT,Integrated DNA technologies) and cloned into the EcoRI/SalI restriction sites of the pPFL-gRNA as described previously ^57^. All plasmid constructs were assembled in *E. coli* prior being transferred to *P. fluorescens* strains SBW25 harboring the aTc-inducible *dcas9* plasmid pPFL-*dcas9*, using standard electroporation techniques.

### 6.4. Analysis of CRISPRi-mediated biofilm and motility phenotype

#### Swarming assay (SW)

The *P. fluorescens* SBW25 strains containing plasmids pPFL-*dcas9* and pPFL-gRNA derivatives were first grown on LB agar plates containing kanamycin (50μg/ml) and gentamycin (10 μg/ml) at 28 °C. Overnight cultures were started from a single fresh colony inoculated in liquid LB medium containing kanamycin and gentamycin and allow to grow at 28 °C with shaking at 220 rpm. Cultures from all strains were then adjusted to OD_600_ = 2 prior to depositing a 1.5 μL drop at the center of a swarming plate (60 mm × 15 mm) containing semi-solid LB-agar (0.6%) complemented with antibiotics and inducer (aTc, 100 ng/ml). Swarming plates were incubated in a growth chamber for 48 h at 25 °C under controlled humidity conditions (70%) before imaging. At least 4 independent swarming surface areas were assessed per strain, with two strains per targeted gene (n=8) using the ImageJ area calculator macrofunction (https://imagej.nih.gov/ij/). The swarming areas were normalized to a reference strain containing a non-targeting pPFL-gRNA plasmid included in each series of experiments. Additionally, a strain expressing a *gacS* (PFLU3777) targeting gRNA was included in each series of tests as a non-swarming control for CRISPR-silencing. The procedure is illustrated in Fig.3B.

#### Biofilm pellicle at air-liquid interface

The formation of biofilm pellicles and imaging were performed as described previously ^57^. Cells carrying pPFL-*dcas9* and pPFL-gRNA derivatives were grown overnight at 28 °C in LB supplemented with kanamycin (50 μg/ml) and gentamycin (10 μg/ml). Overnight cultures were diluted to OD_600_=0.1 in fresh M9-glucose supplemented with the same antibiotics and allowed to grow at 28 °C for 8 hours. All cultures were adjusted to OD_600_=1 and inoculated to OD600 = 0.1 in a 96 wells culture plate containing 200 µl of fresh M9-glucose media supplemented with antibiotics and aTc 100 ng/ml. For each strain, a square of four wells (n=4) were filled. All squares of four wells were separated from each other by empty wells to avoid cross-contamination and to facilitate pellicle collection. Culture plates were incubated at 25 °C and 70% humidity in a growth chamber without shaking for 48 h to allow the complete formation of biofilm pellicles at the liquid-to-air interface. After 48 h, the biofilm pellicles corresponding to four cultures per strain were carefully lifted to the top of the wells by slowly adding M9 media along the side of the wells, such that the convex meniscus reached the top of the wells. Then a cover glass with a 2.5 cm diameter was used to collect the biofilm pellicles by peeling them off the meniscus. The cover glass with the four intact pellicle disks was mounted into an Attofluor microscope chamber (Thermo Fisher Scientific) and the biofilms were dyed by filling with transparent M9 media supplemented with biofilm tracer FM1-43 green biofilm dye (Invitrogen) for 30 min prior to observation by confocal microscopy. Stained biofilms were observed using a spinning disk confocal microscope (Nikon Eclipse Ti-E coupled with CREST X-LightTM confocal imager; objectives Nikon CFI Plan Fluor 10 ×, DIC, 10 × /0.3 NA (WD = 16 mm)). Excitation was performed at 470 nm, and the emission was recorded at 505 nm (green). The procedure is illustrated in Fig. 3C. Images were processed using the Biofilm Analysis XTension package of IMARIS (Bitplane, South Windsor, CT, United States). Biomass, means, and maxima for surface thickness, roughness and surface substratum were determined from at least 4 pellicles and normalized to a reference strain containing a non-targeting pPFL-gRNA plasmid. BioVolume (BV, in µm^3^) is defined as the number of voxels localized inside the isosurface mask. Biomass (BM, in um3/um2) corresponds to the biomass volume divided by the substratum area. BioVolume thickness (maximum MX and Mean MS in µm) was measured from the top of the surface to the base (substratum). The roughness coefficient (R, variance) measures the variability of the localized thicknesses relative to the total mean thickness. A roughness coefficient = 1 suggests a perfectly uniform surface. The larger the variance, the more uneven the surface of the object.

#### Submerged solid-liquid interface biofilms formed on pegs (SBP)

The cell content within submerged biofilms at the solid-liquid interface was monitored using an MBEC Assay® kit, consisting of a 96-well plate covered by a plastic lid holding 96 pegs. Strains carrying pPFL-*dcas9* and pPFL-gRNA derivatives were inoculated at OD600=0.15 in M9 media supplemented with antibiotics and inducer (aTc, 100 ng/ml) to form biofilms at pegs (n=6 pegs per strain, two independent strains per targeted gene). The 96-well setup was incubated at 25 °C and 70% humidity under slow agitation for 48 h. After 48 h, the lid was removed from the culture plate, gently tapped on a paper towel, and allowed to dry. The MBEC lids were then transferred into a 96 well plate filled with 200µL of PBS prior before being sonicated for 30 min in an ultrasonic bath to dislodge and resuspend cells from the biofilms on the pegs. In addition, the liquid cultures (200 µl) which held the pegs were homogenized by sonication. Cell numeration was performed using a flow cytometer (CytoFLEX S; Beckman Coulter). To determine the cell counts at the pegs, 10µl from biofilm resuspension in each well was added to 200uL of PBS. To determine the number of non-adherent cells (remaining in culture) cultures were diluted 1:100 in PSB prior to counting by flow cytometry. The ratio of cells in the biofilm formed pegs-in each well was calculated by dividing the number of cells detected at the pegs by the total number of cells present in the wells (culture+peg). The procedure is illustrated in Fig.3D. It should be noted that the dataset generated by this approach appears to be less robust than those generated for other phenotypes because of the slimy nature of SBW25 biofilms formed at pegs. Despite meticulous manipulation, handling the pegs can result in material loss. This poses a problem when biofilms are formed in limited quantities. However, this assay is particularly well-suited for monitoring enhanced biofilm phenotypes in gene-knockdown experiments.

#### Growth and production of the extra-cellular amyloid fibers (AMF)

The same cultures inoculated for biofilm formation on pegs were also inoculated to determine the production of exopolysaccharides. For each strain, n=5 cultures were inoculated (with two strains per targeted gene) at OD_570_=0.15 in a black clear bottom 96 well plate containing M9 glucose media supplemented with antibiotics, aTc 100 ng/ml, and red fluorescent optotracer (EbbaBiolight 680, Ebba Biotech) for the labeling of extracellular components of the matrix such as amyloids and cellulose secreted in live cultures. Growth (OD_570_) and red fluorescence (Ex. 530 nm, Em. 680 nm) were continuously monitored for 48 h in a multimodal plate reader (Hidex, Lablogic) at 28 °C under continuous agitation. The production of exopolysaccharides was determined by integrating the area under the curve (AUC) after normalization of the fluorescence signal to the OD_570_ (Fig. 3E, Fig. S4).

#### Mitomycin C sensitivity assay

Strains carrying pPFL-*dcas9* and pPFL-gRNA derivatives were grown overnight at 28 °C in LB supplemented with kanamycin (50 μg/ml) and gentamycin (10 μg/ml). Overnight cultures were restarted by diluting to OD_600_=0.01 in fresh LB media supplemented with antibiotics and allowed to grow up to OD_600_=0.2 prior to the addition of aTc (100 ng/ml). The cultures were further grown at 28°C for 5 h to fully induce the expression of *dCas9.* Appropriate dilutions of the cultures were then plated on LB agar plates containing antibiotics and aTc, in the presence or absence of MMC (0.5 µg/ml). The colonies were counted after overnight incubation at 28°C in the dark. Bacterial survival was determined by the ratio of MMC-resistant colonies to the number of colonies in the absence of MMC.

### 6.5. Subcellular localization of FP-labelled c-di-GMP binding proteins and cell morphology

First, the 6His-eGFP coding sequence from the plasmid pJOE7784-1^111^ was replaced with the mNG coding sequence from a synthetic DNA fragment by fragment exchange between EcoRI and HindIII restriction sites, giving rise to the plasmid pJOE7784mod_MCS_V1_mNG (. The genes encoding the CDG proteins DipA (PFLU0458), RbdA (PFLU4308), GcbB (PFLU4600), PFLU5127, PFLU5608, RapA (PFLU2031) MucR (PFLU2753), PFLU1114, AdrA (PFLU3650), Alg44 (PFLU0988), PFLU0007, PFLU0503, PFLU0610, Alg44 (PFLU0988), RimA (PFLU0263), PFLU5210 and PFLU0007 were fused to the mNeonGreen fluorescent protein mNG. For this purpose, the gene coding sequences devoid of their stop codons were PCR-amplified from the SBW25 chromosomal template using oligonucleotide pairs carrying appropriate restriction sites. The amplified DNA fragments were then inserted into the recipient plasmid pJOE7784mod_MCS_V1_mNG between the EcoRI and NotI sites to create a translational fusion with mNG.

Overnight cultures of strains harboring the different plasmid mNG fusion derivatives were used to inoculate fresh LB media at a 1/100 dilution and the cultures were grown to OD600=0,2 prior to adding L-Rhamnose (0,12 mg/ml). At observation time, an aliquot of cells was stained with the amphiphilic styryl red membrane dye FM4-64 (Invitrogen) and laid on the surface of a 1,3% agarose coated glass slide. For single images, time series, and z-stack acquisitions, stained cells expressing a green fluorescent protein fusion were observed using a spinning disk confocal microscope (Nikon Eclipse Ti-E coupled with CREST X-LightTM confocal imager) equipped with a 100x objective lens (CFI Plan Apo Lambda 100x, NA 1.5, WD 130 um). Excitation was performed at 470 nm with emission recorded at 505 nm (green fluorescence) and 520 nm with emission recorded at 630nm (red fluorescence). Image processing was conducted using Fiji ^112^. Cell length analysis was performed using the MicrobeJ plugin ^113^.

### 6.6. Statistical analysis and data availability

All CRISPRi phenotyping experiments were performed at least 4 times for each strain. Unless stated otherwise, all targeted genes were tested twice, either two times independently from the same strain or by testing two independent strains expressing the same gRNA. The mean and standard deviation of the total number of biological replicates for a given gene (n≥8) were reported, and the statistical significance compared with the control strain carrying a non-targeting guide plasmid pPFL_gRNA was assessed using two-tailed Student’s t-tests (Table S5). The statistical significance of annotation and phenotype enrichments was calculated as the Cumulative Hypergeometric Distribution relative to the distribution of annotations in the observed PPI network (Table S3, S6). Principal component analysis (PCA)-based k-means clustering, pairwise Pearson correlation analysis, graphing and illustrations were performed using the OriginPro 2024 data analysis software and visualization package (https://www.originlab.com). The correlation and covariance matrix data are presented in Table S7.

## Author Contributions

MFNG, SF, PL, PN, GB and RW: Conceptualization, Methodology, Investigation. MFNG, PL, RB, and PN Formal analysis, Visualization. GB: Structural modeling analysis. MFNG: Writing original draft. MFNG and PN: Writing - Review & Editing. MFNG, KK and RB: Acquisition of the financial support for the project leading to this publication. All authors have read and approved the final manuscript.

## Acknowledgements

Funding was provided through the Biological Systems Science Division, Office of Biological and Environmental Research, Office of Science, U.S. Dept. of Energy, under Contract DE-AC02-06CH11357. This contribution originates in part from the Environment Sensing and Response Scientific Focus Area (SFA) program and in part from the Small Worlds project at Argonne National Laboratory. Part of the funding contribution also came from INRAE’s internal resources.

## Supplementary information SI-1

- **Rational for the choice of 10 c-di-GMP bait proteins**
- **Supplementary figures**

**Figure S1:** PPI subnetworks organized in functional categories

**Figure S2:** Swarming phenotypic assay

**Figure S3:** Biofilm pellicle phenotypes upon CRISPRi-mediated silencing of genes

**Figure S4:** Effect of gene silencing on the secretion of extracellular amyloid fibers (AMF)

**Figure S5:** Comparative Analysis of CDG Gene Silencing on Biofilm and Swarming Phenotypes

**Figure S6:** Hierarchical clustering of standardized Log2Fold changes across biofilm-related traits for the 37 CDGs

**Figure S7:** Analysis of CDG subnetworks of interaction of (A) DipA (PFLU0458), (B) PFLU5127, (C) BifA (PFLU4858), and (D) Alg44 (PFLU0988) and AwsR (PFLU5210)

**Figure S8:** CGD proteins subcellular localizations

**Figure S9:** Subcellular localization of PFLU5608 during exponential growth

• **Supplementary Tables:**

**Table S1:** c-di-GMP binding proteins domain architecture in SWB25

**Table S2:** c-di-GMP binding proteins y2H screens

**Table S3:** y2H screens functional classes_support to Fig. 1

**Table S4:** Annotations statistical analysis_support to Fig. 5

**Table S5:** Oligonucleotides gene targets

**Table S6:** Biofilm phenotyping data

**Table S7:** Correlation matrix data_support to Fig. 6 and 7

**Supplementary information SI-2:** Structure-prediction benchmark against the Y2H interactome of DipA in Pseudomonas fluorescens SBW25.

**Supplementary Movies:**

Movie S1: Septal localization of PLU5608

Movie S2: Subcellular localization dynamic of RimA

