## Supplementary information SI-1 for "A network perspective on the role of c-di-GMP-associated protein complexes in biofilm formation"

### Supplementary Materials

1. **Rationale for the choice of 10 c-di-GMP bait proteins**

2. **List of Supplementary Figures:**

**Figure S1:** PPI subnetworks organized in functional categories

**Figure S2:** Swarming phenotypic assay

**Figure S3:** Biofilm pellicle phenotypes upon CRISPRi-mediated silencing of genes

**Figure S4:**  Effect of gene silencing on the secretion of extracellular amyloid fibers (AMF)

**Figure S5:** Comparative Analysis of CDG Gene Silencing on Biofilm and Swarming Phenotypes

**Figure S6:** Hierarchical clustering of standardized Log2Fold changes across biofilm-related traits for the 37 CDGs

**Figure S7:** Analysis of CDG subnetworks of interaction of (A) DipA (PFLU0458), (B) PFLU5127, (C) BifA (PFLU4858), and (D) Alg44 (PFLU0988) and AwsR (PFLU5210)

**Figure S8:** CGD proteins subcellular localizations

**Figure S9:** Subcellular localization of PFLU5608 during exponential growth

3. **List of Supplementary Tables:**

**Table S1:** c-di-GMP binding proteins domain architecture in SWB25

**Table S2:** c-di-GMP binding proteins y2H screens

**Table S3:** y2H screens functional classes_support to Fig. 1

**Table S4:** Annotations statistical analysis_support to Fig. 5

**Table S5:** Oligonucleotides gene targets

**Table S6:** Biofilm phenotyping data

**Table S7:** Correlation matrix data_support to Fig. 6 and 7

4. **Supplementary Movies:**

Movie S1: Septal localization of PLU5608

Movie S2: Subcellular localization dynamic of RimA

---------------------------------------------------

#### 1. Rationale for selection of CDG-binding bait proteins:

We performed yeast two-hybrid (Y2H) genomic screens of *P. fluorescens* SBW25 using 10 c-di-GMP-binding (CDG) proteins, PFLU0263, PFLU4858, PFLU1114, PFLU5127, PFLU0458, PFLU4308, PFLU3650, PFLU5210, PFLU0988, and PFLU203 (Table S1). These CDG baits were selected based on i) their involvement in different steps of biofilm formation and dispersal, as reported in the literature, and ii) their involvement in adaptation to environmental changes such as during colonization of plant roots. The c-di-GMP-associated genes PFLU0263, PFLU4858, PFLU1114 and PFLU5127 were found to be upregulated in SBW25 at different stages of rhizosphere colonization [1,2]. PFLU0263 (*rimA*) encodes the phosphodiesterase RimA, which plays a role in the global control of RNA translation [3]. The *rimABK* regulon is upregulated during the early stages of colonization of wheat roots [4]. Its downregulation was found to affect biofilm formation and to increase cell survival against acute exposure to reactive oxygen species [5]. *PFLU4858* encodes the phosphodiesterase BifA, which is involved in biofilm formation in various *Pseudomonas* species. In *P. aeruginosa*, BifA has opposite effects on biofilm and swarming ability [6]. Similarly, in SBW25, the downregulation of *bifA* results in an increased biofilm mass and decreased swarming motility [5]. Consistent with the higher biofilm production, the SBW25 *ΔbifA* mutant also shows higher adhesion to wheat roots [1]. This phenotype has also been observed in *P. putida*, further supporting the involvement of BifA in the surface attachment of cells [1,7]. The protein of unknown function PFLU1114 consists of a conserved EAL domain associated with a degenerate GGDEF domain and a CheY-type receiver domain (REC) characteristic of phosphorylation-activated response regulators. The PFLU1114 gene is part of the plant environmental response regulon [1]. Its downregulation results in increased biofilm mass and structure, suggesting that it carries a phosphodiesterase activity [5]. Similar to *bifA*, deletion of PFLU1114 stimulates attachment at the root surface [1]. It is less functionally characterized, and its mechanism of action during biofilm formation remains to be elucidated. In contrast to BifA and PFLU1114 PDEs, the absence of the DGC gene PFLU5127 shows a reduced ability to bind to root surfaces, suggesting a role at an early stage of biofilm formation [1]. In *P. aeruginosa*, deletion of the PFLU5127-ortholog PA0338 leads to an alteration of the overall structure of biofilms, with a failure to develop particular mushroom-like shapes [8]. The role of PFLU5127 in biofilm formation has not yet been elucidated. The DipA protein, encoded by PFLU0458, also consists of a conserved EAL and degenerate GGDEF domains associated with PAS/GAF clusters. In *P. aeruginosa*, DipA is a membrane-associated PDE involved in cell motility, cell attachment, and biofilm dispersion [9,10]. A *dipA* mutant in this bacterium shows a reduced cell dispersal ability, highlighting its role in the transition from sessile to motile life in the late stage of the biofilm life cycle [8,9]. In SBW25, a *dipA* mutant also exhibits a swarming defect and forms a compact biofilm, suggesting a dispersal deficiency [5]. Similar to DipA, the c-di-GMP-binding protein RdbA encoded by PFLU4308 is a dual GGDEF/EAL domain protein associated with PAS/PAS domains. RdbA has a characterized PDE activity in *P. aeruginosa* and is important for biofilm dispersal [11]. PFLU3650 is a multipass membrane protein containing a single GGDEF domain. PFLU3650 shares a high degree of homology with the transmembrane GGDEF domain protein AdrA of *P. fluorescens* Pf0-1 and P. putida (70% and 62% identity, respectively). AdrA is involved in the regulation of cellulose synthesis by *S. enterica* onto plants [12]. In *P. fluorescens*, AdrA plays a role in flagellar biosynthesis [13,14]. The DCG protein PFLU 5210 is homologous to *P. aeruginosa* DgcN (YfiN), which regulates biofilm integrity in response to oxidative stress [15]. In the plant-associated γ-proteobacterium *Panteoa sp. YR343*, the PFLU5210-ortholog DGC2884 is expressed in the presence of plant roots and is involved in ESP production and biofilm formation [16]. PFLU0988 encodes Alg44, a PilZ-containing protein that regulates the polymerization of alginate, an exopolysaccharide polymer that is widely present in Pseudomonas biofilms [17]. Finally, the protein EAL domain-containing protein PFLU2031 showed significant homology to the phosphodiesterase RapA of *P. fluorescens* Pf0-1, with 87% identical amino acids. In Pf0-1, RapA regulates biofilm formation and breakdown through the release of the LapA adhesin from the cell surface in response to phosphate limitation. However, its role in SBW25, which lacks the Lap system, has yet to be identified [18]. Altogether, these 10 c-di-GMP proteins are involved at various stages of biofilm formation in multiple root-associated Pseudomonas species. They were used as baits in yeast two-hybrid genomic screens of SBW25. A specific design was developed for AdrA to exclude the N-terminal multipass membrane domain.

#### 2. Supplementary Figures

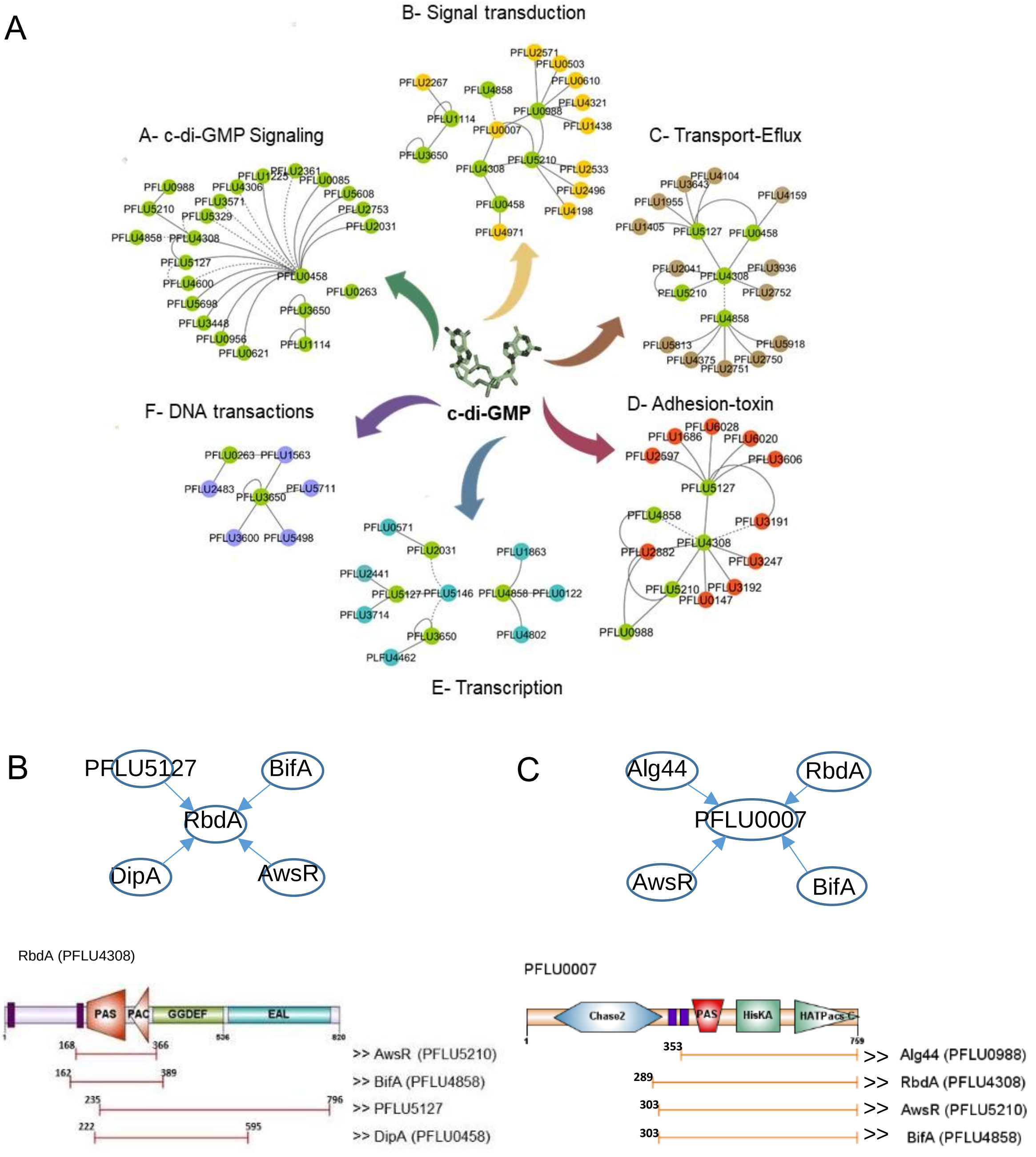


**Figure S1**: PPI subnetworks organized in functional categories. (**A**) PPI-subnetwork connecting c-di-GMP proteins with interacting proteins that belong to the following functional categories. (A) c-di-GMP signaling, (B) signal transduction, (C) transport and efflux, (D) pathogenicity, (E) transcriptional regulation, and (F) DNA repair. Nodes represent proteins, and edges indicate interactions between nodes with good-to-high (solid line) and moderate (dotted line) confidence. (**B and C**) Delineation of minimal interacting domains of RbdA (PFLU4308) and PFLU0007. Functional domains were predicted by SMART (https://smart.embl.de/). GGDEF and EAL domains are shown in green and light blue rectangles, respectively. Accessory domains are indicated in orange pentagon (REC), , blue hexagon (CHASE), red triangle (PAC), red pentagon (PAS), and dark violet segments (transmembrane). **B**- Schematic representation of the RbdA (PFLU4308) interacting landscape. The directionality of the interactions (from bait to prey) is indicated by arrows. Overlapping RbdA domains interacting with AwsR, BifA, DipA and PFLU5127, represented by capped lines, delineate a likely minimal interaction domain. **C**- Schematic representation of the pFLU0007 interacting landscape. The directionality of the interactions is indicated by arrows. Overlapping PFLU0007 domains interacting with AlgAA, RbdA, AwsR, and BifA, illustrated by capped brown lines, delineate a likely minimal interaction domain.

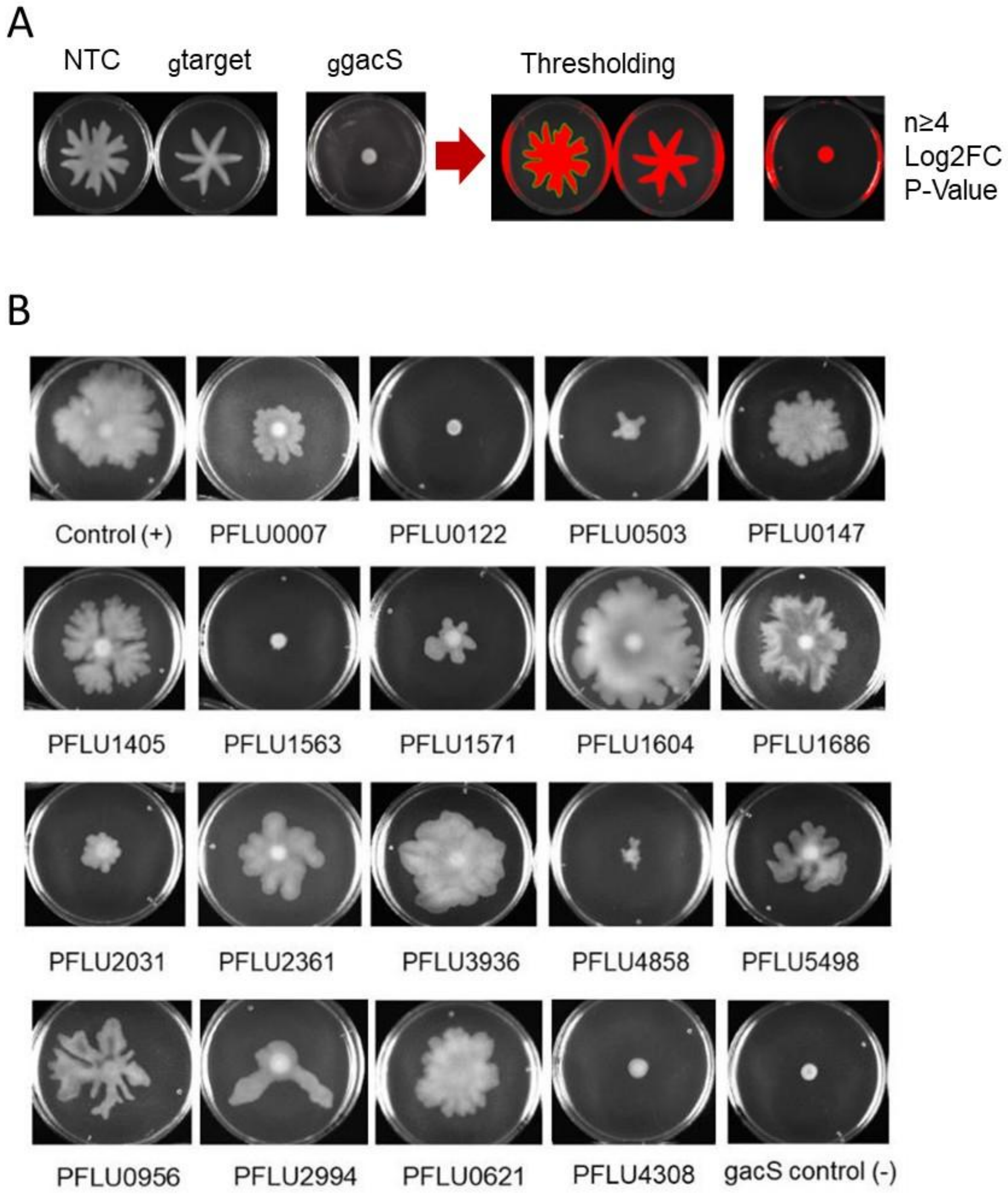


**Figure S2:** *Swarming phenotypic assay*. A- Experimental procedure. Cultures were deposited at the center of semi-solid agar plates containing aTc inducer (100 ng/ml). The plates were incubated at 25°C under controlled humidity for 48 h prior to imaging. Images were processed using ImageJ to determine the swarming area relative to the non-targeting control (NTC) strain expressing dCas9 but targeting no gene. The *gacS* gene has been targeted as an experimental control in all experiments to validate the efficiency of gRNA-mediated silencing. B- Swarming phenotypes exhibited by *P. fluorescens* SBW25 upon CRISPRi-mediated silencing targeting a subset of genes (as indicated).

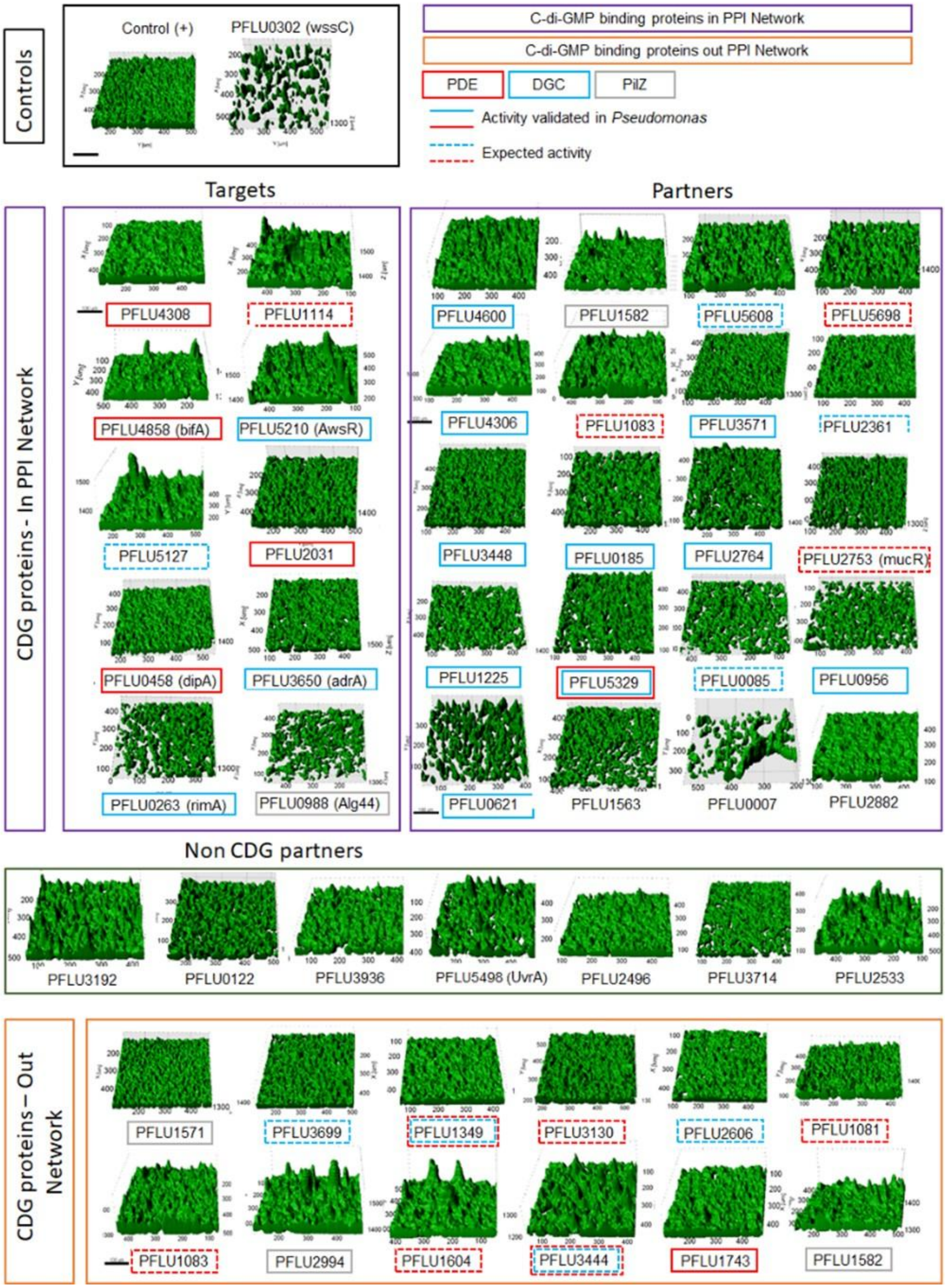


**Figure S3:** *Biofilm pellicle phenotypes upon CRISPRi-mediated silencing of genes*. Cultures were grown at 25°C under controlled humidity for 48 h in LB supplemented with aTc inducer (100 ng/ml). Pellicle were peeled off, mounted on a microscope chamber and stained with FM1-43 prior to observation using confocal microscopy. 3D rendering, visualization, and analysis were performed using Imaris. The targeted genes encoding CDG proteins present in the PPi network (including both bait targets and prey partners), as well as CDG genes not present in the network (Out Network) are indicated. The pellicles formed by a subset of strains targeting genes encoding non-CDGs interacting partners are also shown. A non-targeting control (NTC) strain expressing dCas9 but not targeting any gene along with a strain targeting gene *wssC*, were used as positive (+) and negative controls, respectively.

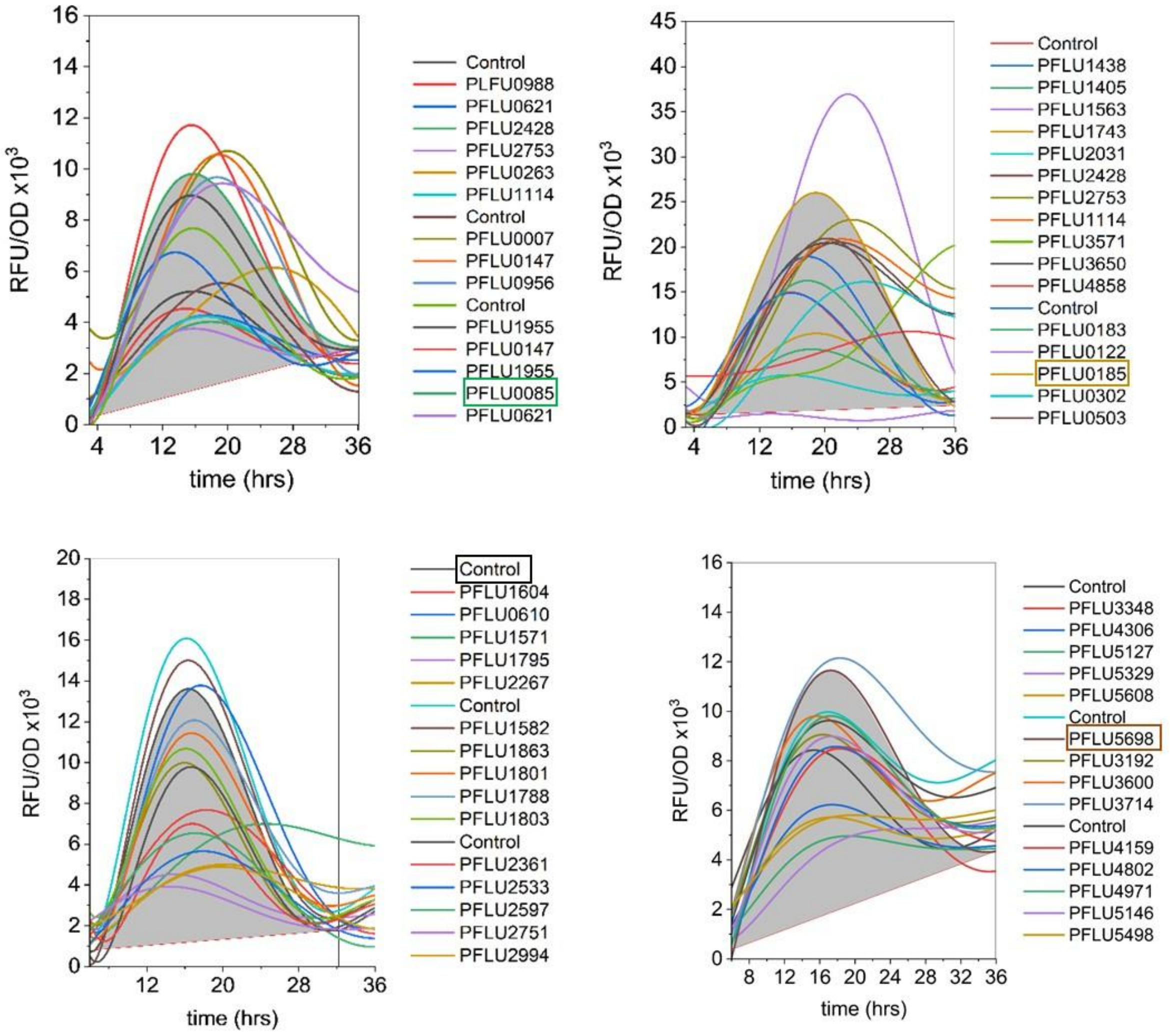


**Figure S4:** *Effect of gene silencing on the secretion of extracellular amyloid fibers (AMF)*. Each culture was performed in the presence of aTc to induce the expression of dCas9 along with red fluorescent optotracer (EbbaBiolight 680) living dye which was used to label secreted amyloid fibers (AMF). The production of AMF was quantified by integrating the area under the curve (AUC) after normalizing the red fluorescence signal against OD (570) (examples of AUC are shown in grey, with the corresponding gene names framed). The control corresponds to strain expressing dCas9 but targeting no gene.

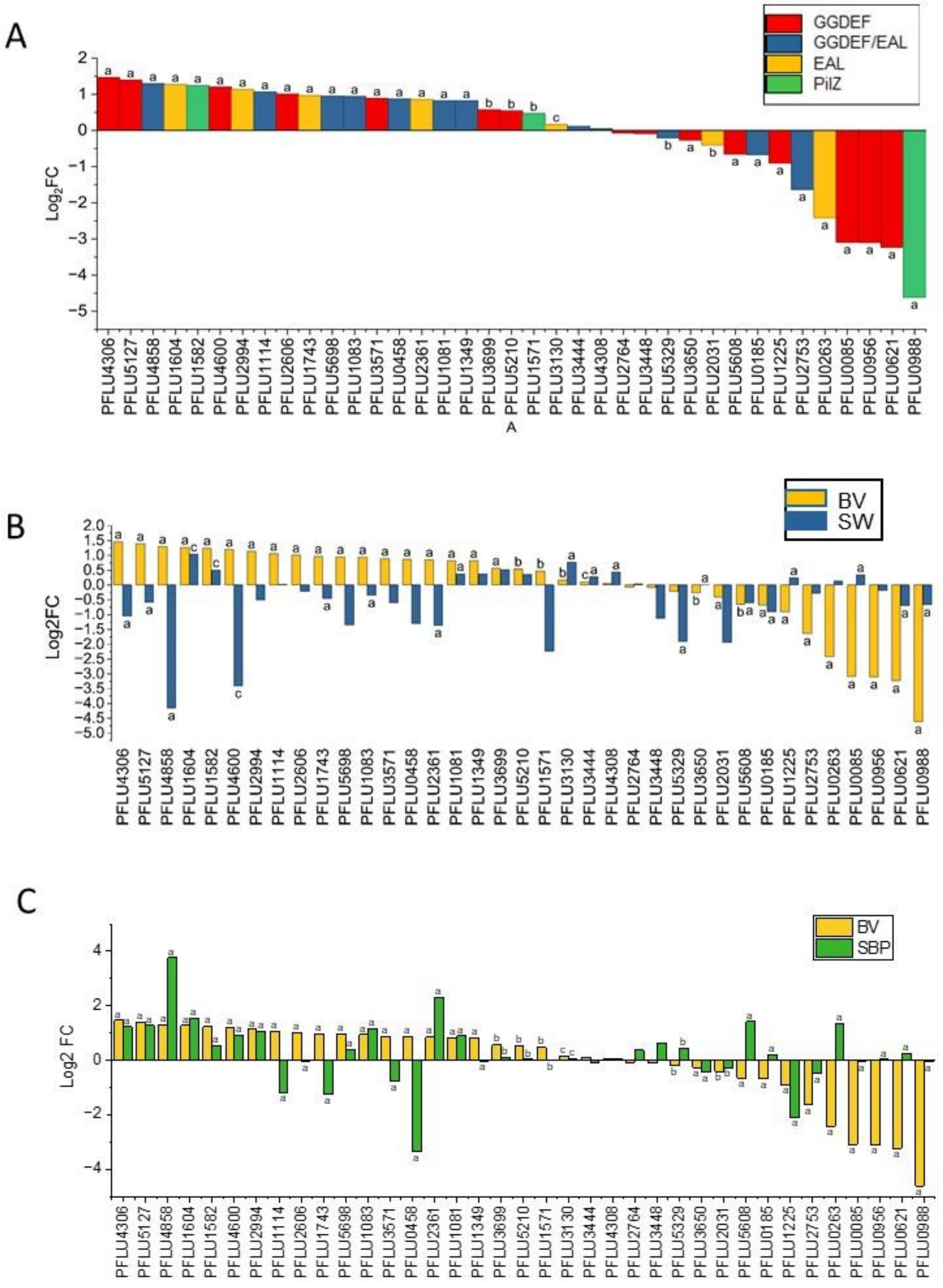


**Figure S5**: *Comparative Analysis of CDG Gene Silencing on Biofilm and Swarming Phenotypes.* (A) The alteration in biofilm volume does not exhibit a strict correlation with the structural organization of CDG protein domains and their predicted activity. The histogram illustrates the Log2fold change in pellicle biovolumes relative to the control strain, which targets no gene. The presence of CDG-domains is indicated by color. (B) Log2fold change in biofilm biovolumes (BV, yellow) is compared to swarming phenotypes (SW, blue). (C) Log2fold change in biofilm biovolumes (BV, yellow) is compared to submerged biofilms at the pegs (SBP, green). Letters denote the levels of statistical significance: (a) p<=0.001, (b) p<=0.01, and (c) p<=0.05.

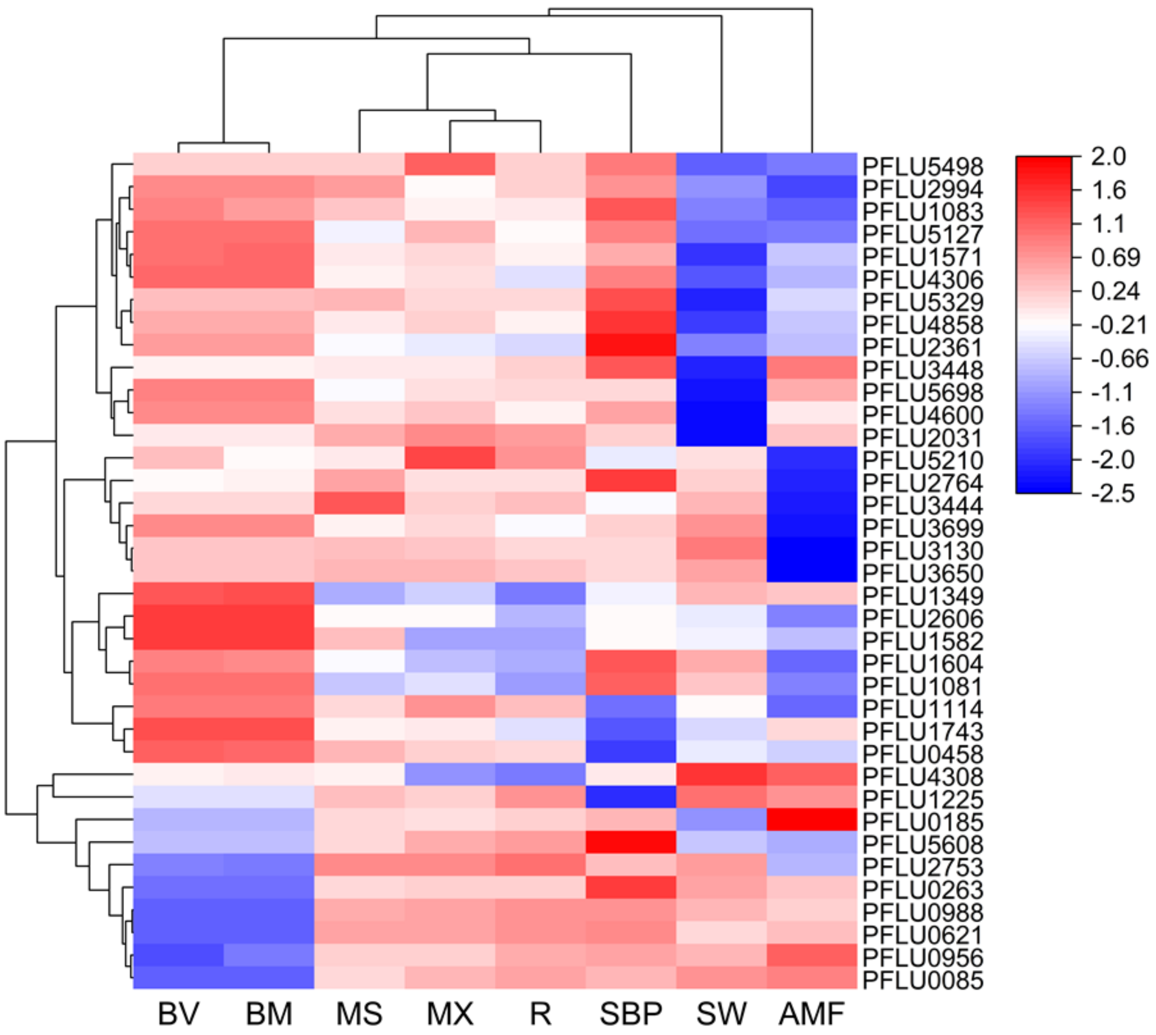


**Figure S6:** Hierarchical clustering of standardized Log2Fold changes across biofilm-related traits for the 37 CDGs. Positive and negative correlations are indicated in shades of red and blue, respectively. Clusters reveal groups of genes with similar phenotypic signatures in biofilm and motility (BV= biovolume, MS= mean thickness, MX= max height, SBP= submerged biofilm, SW= swarming and AMF= amyloid fibers).

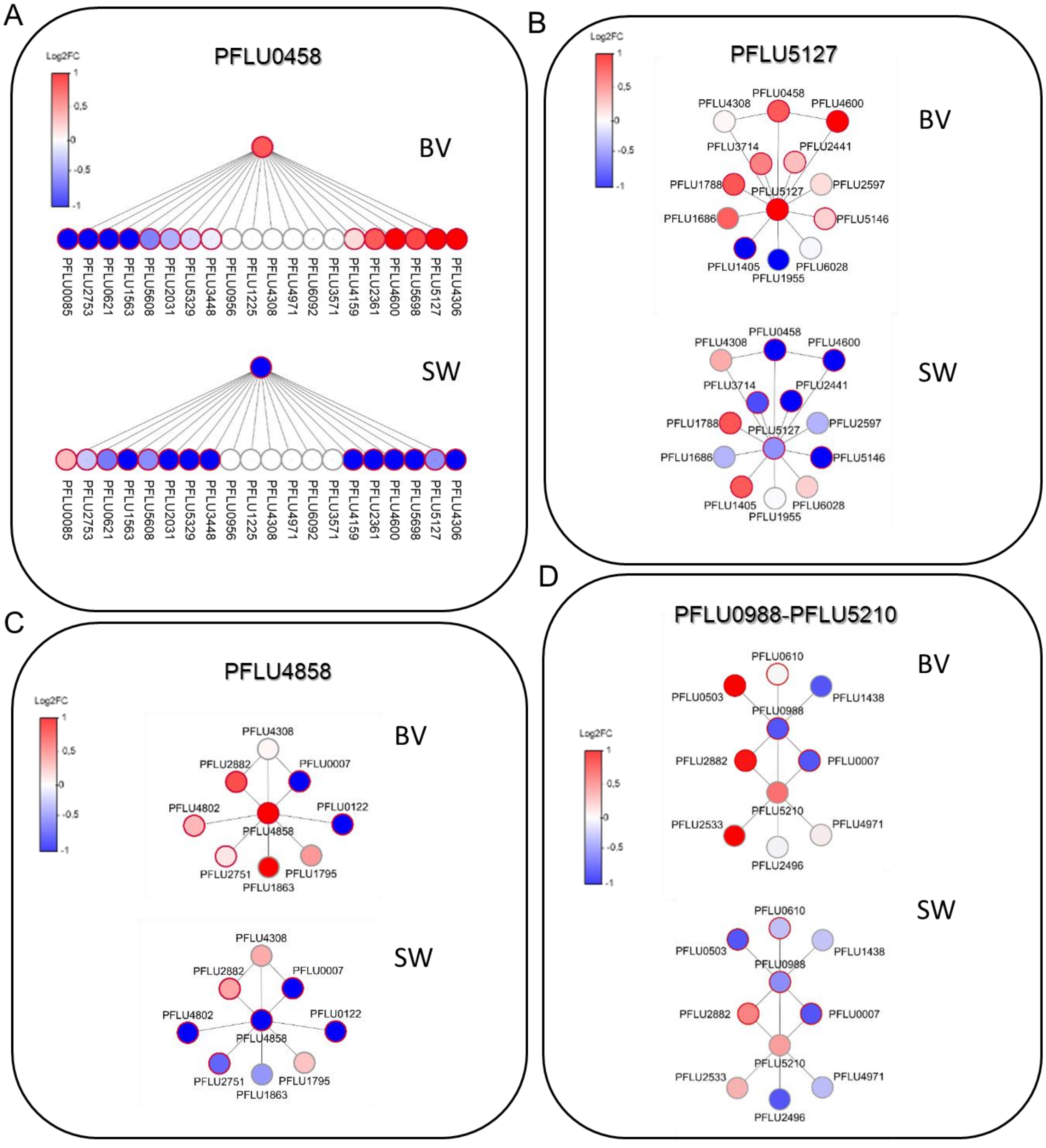


**Figure S7:** Analysis of CDG subnetworks of interaction of (A) DipA (PFLU0458), (B) PFLU5127, (C) BifA (PFLU4858), and (D) Alg44 (PFLU0988) and AwsR (PFLU5210). Proteins are represented as nodes that are connected by edges. Node colors correspond to the Log2 fold change in biofilm volume (BV) and swarming ability (SW) relative to the control strain expressing no guide RNA. Red and blue colors represent enhanced and decreased phenotypes, respectively, with the most intense shades representing a 2 fold increase or decrease, and white indicating no change, as illustrated by the vertical bars. Significance is illustrated by the node border color (red, P<0,05; grey P> 0,05). A node with a non-significant fold change is shown in white.

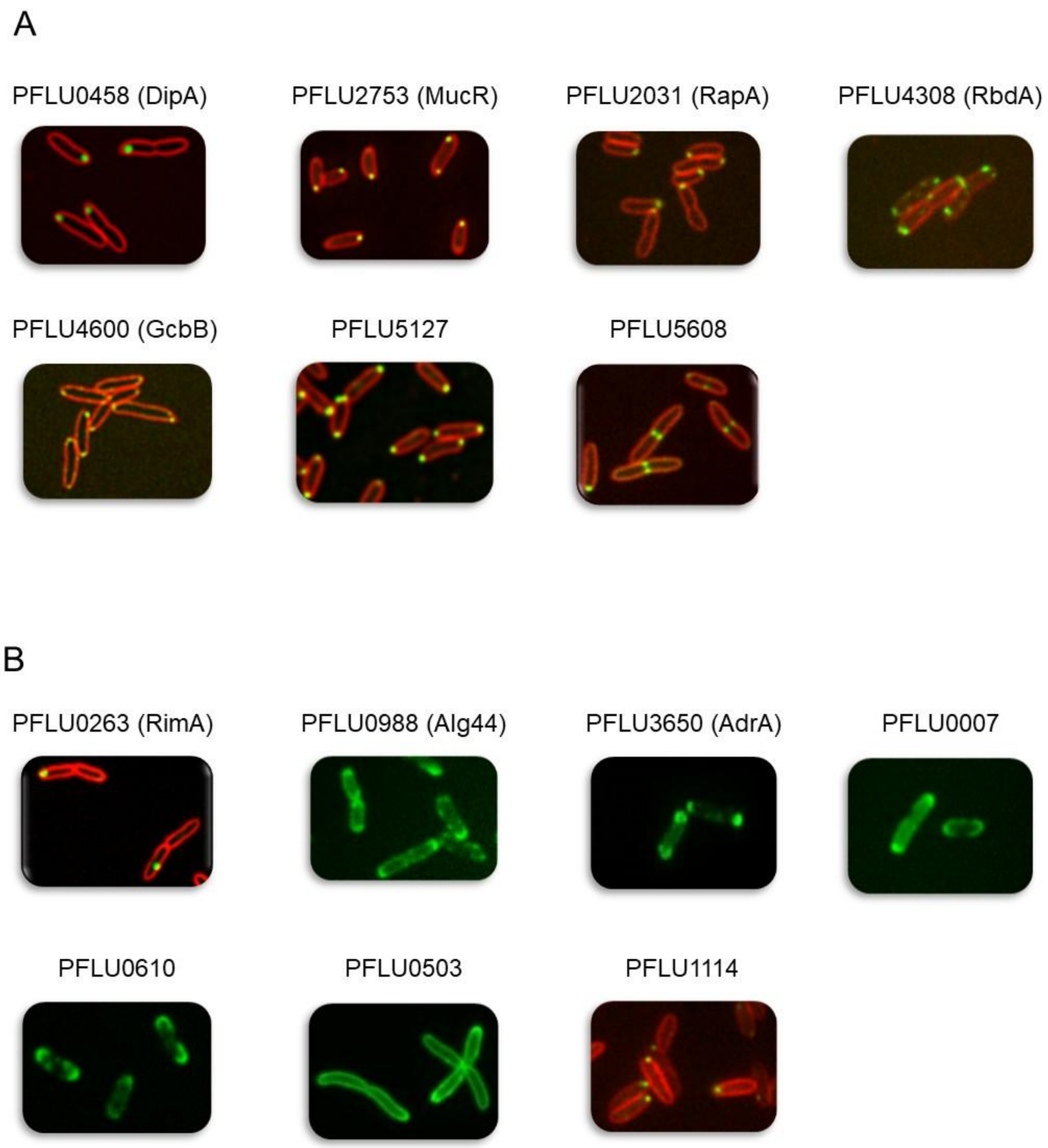


**Figure S8:** *CGD proteins subcellular localizations*. Exponentially growing cells expressing a CDG protein C-terminally fused to mNeonGreen fluorescent protein were observed by spinning disk confocal microscopy. Cell membranes were stained using the lipophilic fluorescent dye FM4-64 (red), except when the fusion protein exhibited a membrane-associated localization (green signal only). (A) GFP-tagged proteins from the DipA subnetwork. Representative images of the discrete localization patterns (green) observed are shown for DipA and six of its interaction partners. (B) Discrete localization profiles for GFP-tagged proteins from other subnetworks.

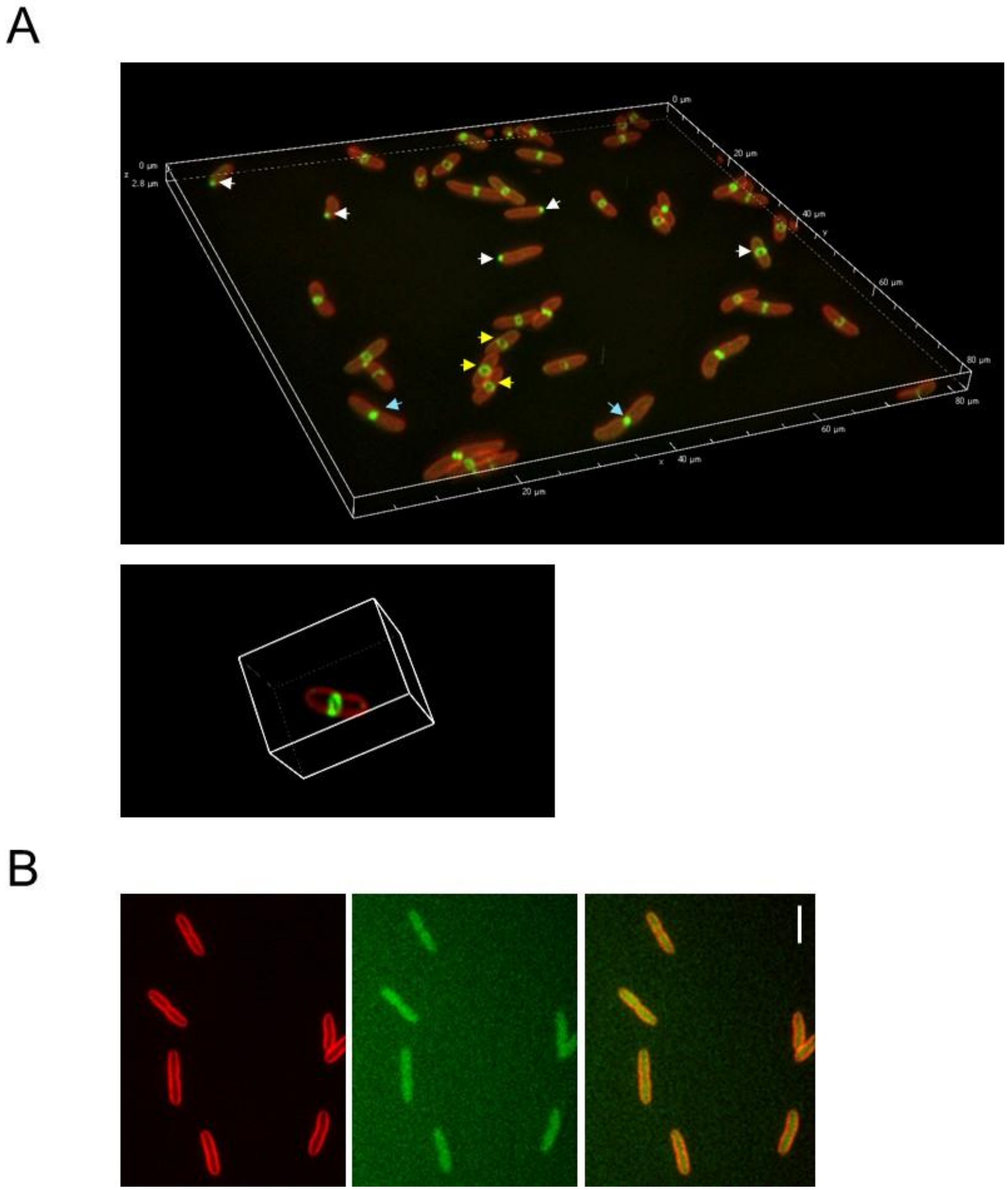


**Figure S9:** *Subcellular localization of PFLU5608 during exponential growth*. (A) Septal localization of PFLU5608 fused to the mNeonGreen fluorescent protein (FP). 3D volume reconstruction of exponentially growing cells expressing PFLU5608-FP (upper panel). Arrows indicate visible ring-like structures (yellow), septal (blue), and polar (white) localizations. An animated view of the 3D volume of a single cell shows a fluorescent ring at the septum (bottom panel) is also available separately. (B) PFLU5210-FP fusion appeared dispersed in the cell with no discrete localization.

1 Little, R. H. *et al.* Differential Regulation of Genes for Cyclic-di-GMP Metabolism Orchestrates Adaptive Changes During Rhizosphere Colonization by Pseudomonas fluorescens. *Frontiers in microbiology* **10**, 1089, doi:10.3389/fmicb.2019.01089 (2019).
2 Silby, M. W. *et al.* Genomic and genetic analyses of diversity and plant interactions of Pseudomonas fluorescens. *Genome Biol* **10**, R51, doi:10.1186/gb-2009-10-5-r51 (2009).
3 Grenga, L. *et al.* Control of mRNA translation by dynamic ribosome modification. *PLoS genetics* **16**, e1008837, doi:10.1371/journal.pgen.1008837 (2020).
4 Little, R. H. *et al.* Adaptive Remodeling of the Bacterial Proteome by Specific Ribosomal Modification Regulates Pseudomonas Infection and Niche Colonisation. *PLoS genetics* **12**, e1005837, doi:10.1371/journal.pgen.1005837 (2016).
5 Noirot-Gros, M.-F., Forrester, S., Malato, G., Larsen, P. E. & Noirot, P. CRISPR interference to interrogate genes that control biofilm formation in Pseudomonas fluorescens. *Scientific Reports* **9**, 1-14 (2019).
6 Kuchma, S. L. *et al.* BifA, a cyclic-Di-GMP phosphodiesterase, inversely regulates biofilm formation and swarming motility by Pseudomonas aeruginosa PA14. *J Bacteriol* **189**, 8165-8178, doi:10.1128/JB.00586-07 (2007).
7 Jimenez-Fernandez, A., Lopez-Sanchez, A., Calero, P. & Govantes, F. The c-di-GMP phosphodiesterase BifA regulates biofilm development in Pseudomonas putida. *Environ Microbiol Rep* **7**, 78-84, doi:10.1111/1758-2229.12153 (2015).
8 Eilers, K. *et al.* Phenotypic and integrated analysis of a comprehensive Pseudomonas aeruginosa PAO1 library of mutants lacking cyclic-di-GMP-related genes. *Frontiers in microbiology* **13**, 949597, doi:10.3389/fmicb.2022.949597 (2022).
9 Roy, A. B., Petrova, O. E. & Sauer, K. The phosphodiesterase DipA (PA5017) is essential for Pseudomonas aeruginosa biofilm dispersion. *J Bacteriol* **194**, 2904-2915, doi:10.1128/JB.05346-11 (2012).
10 Cai, Y. M. *et al.* Differential impact on motility and biofilm dispersal of closely related phosphodiesterases in Pseudomonas aeruginosa. *Sci Rep* **10**, 6232, doi:10.1038/s41598-020-63008-5 (2020).
11 Liu, C. *et al.* Insights into Biofilm Dispersal Regulation from the Crystal Structure of the PAS-GGDEF-EAL Region of RbdA from Pseudomonas aeruginosa. *J Bacteriol* **200**, doi:10.1128/JB.00515-17 (2018).
12 Cowles, K. N., Willis, D. K., Engel, T. N., Jones, J. B. & Barak, J. D. Diguanylate Cyclases AdrA and STM1987 Regulate Salmonella enterica Exopolysaccharide Production during Plant Colonization in an Environment-Dependent Manner. *Appl Environ Microbiol* **82**, 1237-1248, doi:10.1128/AEM.03475-15 (2016).
13 Muriel, C. *et al.* The diguanylate cyclase AdrA regulates flagellar biosynthesis in Pseudomonas fluorescens F113 through SadB. *Sci Rep* **9**, 8096, doi:10.1038/s41598-019-44554-z (2019).
14 Zboralski, A. & Filion, M. Genetic factors involved in rhizosphere colonization by phytobeneficial Pseudomonas spp. *Comput Struct Biotechnol J* **18**, 3539-3554, doi:10.1016/j.csbj.2020.11.025 (2020).
15 Katharios-Lanwermeyer, S., Koval, S. A., Barrack, K. E. & O'Toole, G. A. The Diguanylate Cyclase YfiN of Pseudomonas aeruginosa Regulates Biofilm Maintenance in Response to Peroxide. *J Bacteriol* **204**, e0039621, doi:10.1128/JB.00396-21 (2022).
16 Bible, A. N., Chang, M. & Morrell-Falvey, J. L. Identification of a diguanylate cyclase expressed in the presence of plants and its application for discovering candidate gene products involved in plant colonization by Pantoea sp. YR343. *PLoS One* **16**, e0248607, doi:10.1371/journal.pone.0248607 (2021).
17 Merighi, M., Lee, V. T., Hyodo, M., Hayakawa, Y. & Lory, S. The second messenger bis-(3'-5')-cyclic-GMP and its PilZ domain-containing receptor Alg44 are required for alginate biosynthesis in Pseudomonas aeruginosa. *Molecular microbiology* **65**, 876-895, doi:10.1111/j.1365-2958.2007.05817.x (2007).
18 Newell, P. D., Boyd, C. D., Sondermann, H. & O'Toole, G. A. A c-di-GMP effector system controls cell adhesion by inside-out signaling and surface protein cleavage. *PLoS Biol* **9**, e1000587, doi:10.1371/journal.pbio.1000587 (2011).
