## Supplementary information SI-2 for "A network perspective on the role of c-di-GMP-associated protein complexes in biofilm formation"

**Benchmarking of structure prediction against the yeast two-hybrid interactome of PDE PFLU0458.**

To assess whether AI-based structure prediction can identify experimentally observed protein–protein interactions, we analyzed a yeast two-hybrid (Y2H) interaction dataset generated using the c-di-GMP signaling protein PFLU0458 from Pseudomonas fluorescens SBW25 as bait. Protein complexes were predicted with Boltz-2 and evaluated using the structure-based binding affinity predictor PRODIGY. Predicted interactions were benchmarked against three annotated reference sets consisting of Y2H-positive interactors, sequence-distant proteins, and putative negative interactors, along with monomeric controls.

### Methods

#### Experimental interaction dataset.

Protein–protein interactions identified for the bait protein PFLU0458 by yeast two-hybrid (Y2H) screening were used to define the benchmark dataset. For benchmarking, prey proteins were classified into three reference sets: Y2H-positive interactors (n = 14), putative negative interactors with GGDEF domain (n = 5) selected from the c-di-GMP signaling protein family, and sequence-distant proteins (n = 3) exhibiting detectable sequence similarity to PFLU0458 but not identified as interactors in the Y2H screen. In addition, PFLU0458 and its highest-confidence interacting partner, PFLU0956, were modeled individually as monomers to assess the quality of the predicted protein structures.

#### Complex structure prediction

Protein complexes were predicted for each bait–prey pair using Boltz-2 via the Neurosnap platform (accessed July 2026). Each bait and prey sequence was modeled as a 1:1 heterodimer, and five independently seeded models (rank_1–rank_5) were generated using the default multiple sequence alignment (MSA)-based pipeline. For each prediction, the platform reported confidence metrics together with predicted aligned error (PAE) and predicted distance error (PDE) matrices. At the time of analysis, structural predictions were unavailable for two of the five putative negative proteins; consequently, the final benchmark dataset comprised 20 predicted complexes (14 Y2H-positive, 3 putative negative, and 3 sequence-distant pairs) together with two monomer models.

#### Confidence and interface score extraction

Confidence metrics were extracted from each Boltz-2 prediction archive using an in-house Python script (parse.neurosnap.py). For each of the five predicted models, the following metrics were collected: interface predicted TM-score (ipTM), chain-pair ipTM, global predicted TM-score (pTM), interface predicted distance error (complex_iPDE), interface pLDDT (complex_iPLDDT), pDockQ, pDockQ2, interface predicted score of aligned error (ipSAE; calculated at the chain, domain, and residue levels using multiple $d_0$ values), and the Local Interaction Score (LIS). For each bait–prey pair, best (rank 1), maximum, mean, and standard deviation values were calculated across the five models. The resulting datasets are provided as a per-pair summary matrix (neurosnap.matrix.tsv) and a per-model results table (neurosnap.permodel.tsv).

#### Binding affinity estimation

For the Y2H-positive complex that exhibited clear structural support (see Results), binding affinity was estimated using PRODIGY (prodigy-prot v2.4.0). Binding affinities were calculated for each of the five Boltz-2 models at 25 °C using default parameters, including a 5.5 Å intermolecular contact cutoff and solvent accessibility calculated with FreeSASA v2.2.1. Biopython v1.85 was used for structure processing.

### Results

#### Interface confidence scores do not recapitulate the Y2H interaction map

Across the 20 predicted bait–prey complexes, **pDockQ** and **pDockQ2** remained at their minimum values (0.0183 and 0.0073, respectively), indicating that Boltz-2 did not predict a confident, contact-rich interface for any complex. Consequently, neither metric provided discriminatory power for distinguishing Y2H-positive interactions from the reference datasets (Table 1). The remaining interface metrics, including **ipTM**, **ipSAE**, **LIS**, and **complex_iPDE**, also failed to separate Y2H-positive interactors from putative negative or sequence-distant proteins. Instead, the highest-scoring predictions were obtained for a sequence-distant protein (**PFLU0263**; ipTM = 0.74) and a putative negative protein (**PFLU0185**; complex_iPDE = 5.0 Å, ipSAE = 0.39), both of which ranked above nearly all Y2H-positive complexes. Consistent with these observations, the Y2H-positive set exhibited a median best-model ipSAE value of **0.000**, indicating that more than half of the experimentally identified interactors showed no detectable interface signal, compared with median values of **0.267** and **0.274** for the putative negative and sequence-distant reference sets, respectively. Applying commonly used confidence thresholds (ipTM > 0.8 and pDockQ > 0.23) failed to identify any of the 14 Y2H-positive interactions as bona fide complexes. This finding is consistent with the known challenges of predicting transient, regulated, or domain-mediated protein–protein interactions, which are thought to predominate within c-di-GMP signaling networks. These results indicate that standard confidence thresholds developed for stable protein complexes are not directly applicable to this class of interactions.

**Table 1.** Interface scores by reference category (best model per pair; median / best).

| **Category** | **n** | **ipTM (Median / Maximum)** | **ipSAE (d0res) (Median / Maximum)** | **LIS (Median / Maximum)** | **Complex iPDE (Å) [1] (Median / Minimum)** | **pDockQ** |
|---|---|---|---|---|---|---|
| Y2H-positive | 14 | 0.452 / 0.554 | 0.000 / 0.405 | 0.025 / 0.217 | 13.2 / 5.6 | 0.0183 (floor) |
| Putative negative | 3 | 0.447 / 0.580 | 0.267 / 0.392 | 0.170 / 0.213 | 10.7 / 5.0 | 0.0183 (floor) |
| Sequence-distant | 3 | 0.572 / 0.736 | 0.274 / 0.384 | 0.143 / 0.229 | 8.5 / 8.1 | 0.0183 (floor) |

[1] Complex iPDE is an error term - lower is better; the second value is the minimum.

#### A single structurally-supported positive interaction

Among the Y2H-positive interactions, the **PFLU0458–PFLU5698** complex exhibited the strongest structural support. The highest-ranked model achieved an ipTM of **0.55**, an ipSAE of **0.41** (the highest observed among all predicted complexes), an LIS of **0.22**, and a complex_iPDE of **5.6 Å**. Binding affinity estimation using PRODIGY across the five Boltz-2 models predicted a consistent, well-defined interface comprising an average of **451 intermolecular contacts** (range, 391–486) and a mean binding free energy of **ΔG = −37.4 ± 3.1 kcal·mol$^{-1}$** (range, −32.3 to −41.4; Table 2). The highest-ranked model contained **477 intermolecular contacts**, including 125 charged–apolar, 108 apolar–apolar, 98 apolar–polar, 64 charged–polar, 57 charged–charged, and 25 polar–polar interactions. The corresponding non-interacting surface consisted of **39.0% apolar** and **29.2% charged** residues.

**Table 2.** PRODIGY per-model results for PFLU0458–PFLU5698 (25 °C).

| Model | Intermolecular Contacts | Predicted Binding Free Energy, ΔG (kcal·mol$^{-1}$) | Predicted Dissociation Constant, K<sub>d</sub> (M) |
|---|---|---|---|
| Rank 1 | 477 | −39.8 | 5.9 × 10<sup>−30</sup> |
| Rank 2 | 449 | −36.3 | 2.5 × 10<sup>−27</sup> |
| Rank 3 | 486 | −41.4 | 4.0 × 10<sup>−31</sup> |
| Rank 4 | 391 | −32.3 | 2.0 × 10<sup>−24</sup> |
| Rank 5 | 452 | −37.2 | 5.6 × 10<sup>−28</sup> |
| **Mean** | **451** | **−37.4** | - |

**Important caveat.** The predicted binding affinities should be interpreted with caution because they were obtained by extrapolating beyond the validated range of the PRODIGY model. The protein complexes used for PRODIGY training and validation typically contained substantially fewer intermolecular contacts (tens rather than hundreds) and exhibited experimentally determined binding free energies of approximately −5 to −18 kcal·mol$^{-1}$. In contrast, the PFLU0458-PFLU5698 models contained approximately 450 intermolecular contacts, forcing the underlying linear IC-NIS model to extrapolate beyond its validated domain. Consequently, the predicted ΔG and sub-picomolar $K_d$ values are unlikely to be quantitatively reliable. Instead, these results should be interpreted qualitatively, indicating that Boltz-2 consistently predicts a large protein–protein interface and that PRODIGY estimates this interface to be energetically favorable if the predicted structure is accurate. Despite these observations, pDockQ remained at its minimum value, highlighting the uncertainty of the predicted interaction. Therefore, independent experimental validation, such as isothermal titration calorimetry (ITC), surface plasmon resonance (SPR), microscale thermophoresis (MST), or interface-directed mutagenesis, will be required to establish whether PFLU0458-PFLU5698 forms a bona fide protein complex.

#### PRODIGY analysis of representative negative and sequence-distant complexes

To evaluate whether structure-based binding affinity estimates could distinguish experimentally supported interactions from non-interacting complexes, PRODIGY was applied to the highest-scoring representative from each reference class. These included the Y2H-positive complex **PFLU0458-PFLU5698**, the putative negative complex **PFLU0458-PFLU0185** (lowest complex_iPDE in the dataset; 5.0 Å), and the sequence-distant complex

**PFLU0458-PFLU0263** (highest ipTM in the dataset; 0.74). Binding affinities were calculated for all five Boltz-2 models of each complex.

**Table 3.** PRODIGY binding affinity predictions for representative complexes from each benchmark class (mean ± SD across five Boltz-2 models at 25 °C).

| **Complex** | **Reference class** | **Mean intermolecular contacts** | **Mean ΔG (kcal·mol$^{-1}$)** | **ΔG range (five models)** |
|---|---|---|---|---|
| PFLU0458–PFLU5698 | Y2H-positive | 451 | −37.4 ± 3.1 | −32.3 to −41.4 |
| PFLU0458–PFLU0185 | Putative negative | 323 | −25.3 ± 1.3 | −22.7 to −26.1 |
| PFLU0458–PFLU0263 | Sequence-distant | 131 | −14.1 ± 3.3 | −9.5 to −18.4 |

PRODIGY predictions did not improve discrimination between experimentally supported and unsupported interactions. Instead, the predicted binding free energies closely tracked the size of the modeled protein–protein interface, as reflected by the number of intermolecular contacts, rather than the Y2H interaction status. Although **PFLU0458–PFLU0185** was classified as a putative negative interactor, it was predicted to form an extensive 323-contact interface with a mean binding free energy of −25.3 kcal·mol$^{-1}$, values that would ordinarily be interpreted as indicative of a stable complex. In contrast, the sequence-distant complex (**PFLU0458–PFLU0263**) contained only 131 intermolecular contacts and exhibited substantially weaker predicted binding affinities (ΔG = −14.1 ± 3.3 kcal·mol$^{-1}$). Notably, this was the only representative complex whose predicted contact numbers and binding free energies fell within the validated operating range of the PRODIGY model. Together with the interface confidence analysis, these findings indicate that neither Boltz-2 confidence metrics nor PRODIGY binding affinity predictions accurately recapitulate the Y2H interaction map for **PFLU0458**.

Per-model PRODIGY results are provided in PFLU0458_PFLU5698.prodigy.tsv, Neg_PFLU0458_PFLU0185.prodigy.tsv, and Distant_PFLU0458_PFLU0263.prodigy.tsv.

### Discussion

The present analysis represents a proof-of-principle evaluation of whether AI-based protein complex prediction can recover experimentally determined protein–protein interactions identified by yeast two-hybrid screening. Although Boltz-2 successfully generated structural models for all bait–prey combinations, neither its interface confidence metrics nor subsequent binding affinity estimation with PRODIGY reliably distinguished Y2H-positive interactions from putative negative or sequence-distant proteins. These findings suggest that current structure-based confidence metrics are insufficient for identifying transient signaling interactions within c-di-GMP regulatory networks.

Several factors likely contribute to this outcome. First, the benchmark was intentionally limited to a single bait protein (PFLU0458), five independently generated models per bait–prey pair, and a relatively small set of experimentally supported and control proteins. Consequently, the present results should be regarded as qualitative rather than statistically definitive. Expanding the analysis to additional bait proteins and larger interaction datasets will be necessary to determine whether these observations generalize across c-di-GMP signaling systems.

Second, many signaling proteins are expected to participate in transient, weak, or condition-dependent interactions that differ fundamentally from the stable complexes used to train most current structure prediction and interface-scoring methods. Similar to AlphaFold-Multimer, Boltz-2 can generate extensive protein–protein interfaces even for complexes that may not exist under physiological conditions. The large intermolecular contact networks predicted for several putative negative complexes illustrate this limitation and emphasize that extensive structural interfaces alone should not be interpreted as evidence of biological interaction.

The limitations of the subsequent PRODIGY analysis should also be considered. PRODIGY assumes that the input structure represents a correct bound complex and estimates binding affinity directly from the predicted interface geometry. Consequently, any inaccuracies in the structural model propagate directly into the predicted binding free energies. Furthermore, the exceptionally large interfaces predicted for several Boltz-2 models extend well beyond the calibration range of PRODIGY, making the resulting ΔG and Kd values unsuitable for quantitative interpretation. Instead, these predictions should be viewed primarily as qualitative measures of interface consistency rather than estimates of true binding affinity.

Despite these limitations, the combined Boltz-2 and PRODIGY analyses identified one interaction, **PFLU0458-PFLU5698**, that consistently exhibited stronger structural support than any other Y2H-positive complex. While this prediction requires independent experimental validation using orthogonal approaches such as isothermal titration calorimetry, surface plasmon resonance, microscale thermophoresis, or mutational analysis, it demonstrates that AI-based structure prediction can serve as a useful prioritization tool for selecting candidate interactions for detailed biochemical characterization. Future improvements in modeling transient protein complexes, together with the incorporation of experimental constraints and larger benchmark datasets, may substantially improve the ability of AI-based methods to reconstruct bacterial signaling networks.

### Software and data availability

Boltz-2 structure predictions were generated using the Neurosnap platform (accessed July 2026), with five independently seeded models produced for each bait–prey complex. Prediction confidence metrics were extracted using an in-house Python script neurosnap.py (available upon request). Processed results are provided as the per-pair summary matrix (neurosnap.matrix.tsv), the per-model results table (neurosnap.permodel.tsv), and the corresponding PRODIGY output files (PFLU0458_PFLU5698.prodigy.tsv, Neg_PFLU0458_PFLU0185.prodigy.tsv, and Distant_PFLU0458_PFLU0263.prodigy.tsv). Binding affinity estimates were calculated using PRODIGY (prodigy-prot v2.4.0) with FreeSASA v2.2.1 and Biopython v1.85.
